# Fusion-negative Rhabdomyosarcoma 3D-organoids as an innovative model to predict resistance to cell death inducers

**DOI:** 10.1101/2022.09.06.506756

**Authors:** Clara Savary, Paul Huchedé, Léa Luciana, Arthur Tourbez, Clémence Deligne, Cécile Picard, Thomas Diot, Claire Coquet, Nina Meynard, Marion Le Grand, Laurie Tonon, Nicolas Gadot, Cyril Degletagne, Sophie Léon, Valéry Attignon, Alexandra Bomane, Isabelle Rochet, Kevin Müller, Virginie Mournetas, Christophe Bergeron, Paul Rinaudo, Aurélie Dutour, Martine Cordier-Bussat, Frédérique Dijoud, Nadège Corradini, Delphine Maucort-Boulch, Eddy Pasquier, Jean-Yves Blay, Marie Castets, Laura Broutier

## Abstract

Rhabdomyosarcoma (RMS) is the main form of soft-tissue sarcoma in children and adolescents. For 20 years, and despite international clinical trials, its cure rate has not really improved, and remains stuck at 20% in case of relapse. The definition of new effective therapeutic combinations is hampered by the lack of reliable models, which complicate the transposition of promising results obtained in pre-clinical studies into efficient solutions for young patients. Inter-patient heterogeneity, particularly in the so-called fusion-negative group (FNRMS), adds an additional level of difficulty in optimizing the clinical management of children and adolescents with RMS.

Here, we describe an original 3D-organoid model derived from relapsed FNRMS and show that it finely mimics the characteristics of the original tumor, including inter- and intra-tumoral heterogeneity. Moreover, we have established the proof-of-concept of their preclinical potential by re-evaluating the therapeutic opportunities of targeting apoptosis in FNRMS from a streamlined approach based on the exploitation of bulk and single-cell omics data.

## Introduction

Triggering tumor cell elimination through activation of death signaling pathways conceptually appears to be one of the most direct methods to cure patients with cancer^1^. Among cell death mechanisms, induction of tumor cells’ apoptosis seemed particularly promising, given the finely regulated nature of this process at the molecular level. Apoptosis is one of the major forms of programmed cell death by which a supernumerary, ectopic or abnormal cell triggers its own elimination^2^. Reciprocally, resistance to apoptosis is considered as one of the first 6 features described as acquired by cells during tumor initiation and progression^3^. Then, the development of drugs capable of restoring the execution of apoptosis in tumor cells has emerged as a significant therapeutic lever^1^. However, despite promising results *in vitro* and in preclinical trials, these therapies have so far been disappointing in patients^4,5^. One of the reasons for these results could be the lack of appropriate models to select therapeutic combinations sufficient to efficiently trigger apoptotic pathways in tumor cells, taking into account the intrinsic complexity of these signaling cascades and of the tumor ecosystem^6–8^.

This pitfall could notably apply to rhabdomyosarcoma (RMS). RMS are rare cancers, representing 5% of pediatric solid tumors and affecting 4 children/adolescents per million^9^. RMS are characterized by their similarities to embryonic muscle tissue, including the expression of specific markers such as Desmin, MYOD1, or Myogenin, which are routinely used in their differential diagnosis^10,11^. Apart from this single common denominator, RMS are a heterogeneous group of cancers. Two main subclasses of RMS have been defined based on histological features in the pediatric population. More than 70% of all RMS falls into the embryonal RMS subclass (ERMS) and 20% into the alveolar rhabdomyosarcoma (ARMS) one. In contrast to ERMS, ARMS are more common in older children and young adolescents^12^. Considerable advances in understanding the molecular etiology of RMS has resulted from the identification of pathognomonic chromosomal translocations associated with 85% of ARMS, which lead to the expression of oncogenic fusion transcription factors, PAX3-FOXO1 or PAX7-FOXO1^13^. However, the molecular bases of the group gathering ERMS and ARMS lacking PAX-FOXO1 translocation are more complex. Indeed, these so-called fusion-negative RMS (FNRMS) could result from recurrent single nucleotide variations in a number of well-characterized oncogenes such as *HRAS, NRAS, KRAS, ALK, FGFR4, PIK3CA, FBXW7, NF1, TP53, CTNNB1*, or *BCOR*^14^, while some sequenced tumors do not have identifiable driver mutations^14^. Fusion-positive RMS (FPRMS) tumors are generally considered as high-risk, notably due to their higher propensity to metastasize^15,16^, but the situation is more complicated in the heterogeneous FNRMS entity. Indeed, no reliable molecular alteration that could help clinicians to adapt treatment intensity is available so far^16^. Treatment commonly combines an aggressive neo-adjuvant chemotherapy (Vincristine, Dactinomycin, and Cyclophosphamide/Ifosphamide) with secondary surgery, associated more often with radiotherapy, and maintenance chemotherapy for high-risk patients. Considering the long-term side effects of such intensive multimodal regimens in early life^17^, new molecular tools are required to improve FNRMS patients’ stratification in risk groups. Irradiation of children, especially those less than 36 months, is notably challenging given the risk of sequelae and associated morbidity and must be reserved for the most aggressive cases. Moreover, the 5 year-survival rate ranges from 60 to 80% for patients with localized tumors, with only a slight improvement in the prognosis over the last 20 years^16,18^. Only 20% of RMS that have relapsed or with metastases at the time of diagnosis can be cured^16^.

Targeting apoptosis has been considered as a putative therapeutic lever in RMS^6,19–23^. Indeed, a shift towards cell survival resulting notably from the aberrant activation of the RAS/PI3K pathway is one of the main known oncogenic hallmarks of FNRMS^11^. Alterations in the expression of some apoptotic effectors have been linked to clinical outcome^24,25^ and several studies have evaluated the therapeutic potential of targeting apoptosis, mostly *in vitro* on cell lines grown as monolayers and *in vivo* in xenograft experiments^19^. However, the potential for clinical translation of these data is hampered by the anticipation of the emergence of resistance, which has been observed in clinical trials in other cancers^6,26^.

We propose here to reconsider the therapeutic potential of targeting apoptosis in RMS, especially the fusion-negative ones, starting from an integrative transcriptomic analysis of apoptotic pathways. By establishing a comprehensive cartography of these pathways, we show here that FNRMS can be considered as primed-for-death, and highlight the potential clinical value of this death pathway for risk stratification. More importantly, we have developed an original 3D patient-derived organoid model, which recapitulates aggressive FNRMS tumors’ histological and molecular characteristics. By combining bioinformatic analyses of bulk and single-cell transcriptome data and using these 3D-innovative models, we prove that it is possible to improve the efficacy of therapeutic strategies focused on apoptosis by a streamlined approach combining the precise and simultaneous targeting of blocking points and intra-tumor heterogeneity.

## Results

### Apoptosis is committed and has a prognostic value in FNRMS based on bulk tumor transcriptomic data integration

Until now, apoptotic blockage in RMS has been evaluated mostly at discrete levels, without trying to reconstruct the global state of functionality of associated pathways. Starting from bulk transcriptomic profiling of independent RMS cohorts^13,14,27^ (Extended Data Fig. 1), we first analyzed RMS apoptotic signature heterogeneity using dimensionality reduction methods^28^. Plots by uniform manifold approximation and projection (UMAP) and principal component analysis (PCA) demonstrated separated clusters for RMS (in yellow) and healthy control muscles (in blue) in two independent cohorts, solely based on the expression of an apoptotic signature^29^ (Fig. 1a, Extended Data Fig. 2a and Supplementary Table 1). Similarly, FNRMS and FPRMS cluster largely separate based on this apoptotic signature in plots generated by UMAP and PCA (Fig. 1b and Extended Data Fig. 2b-c). To explore the biological significance of this clustering, we used causal inference approaches to define the apoptotic cascades’ activation level status based on the expression profile of their effectors^30^. Activation of a transcriptional regulatory network predicting engagement notably of the Death Receptors (DR) extrinsic signaling was identified by Ingenuity Pathway Analysis (IPA) in RMS tumors compared to normal muscle samples in cohort 1 (p-value < 0.0001, z-score = 0.60) and cohort 2 (p-value < 0.0001, z-score = 1.94; Fig. 1c). Z-scores positivity of apoptotic pathways was also significantly higher in FNRMS compared to FPRMS in cohort 3 (p-value < 0.0001, z-score = 1.94; Fig. 1d) and cohort 4 (p-value < 0.0001, z-score = 1.41; Extended Data Fig. 2d). In addition, we used the PROGENy method to infer downstream apoptotic-response footprint from perturbation-response genes indicative of the DR TRAIL activity^31^. As shown in Fig. 1e, TRAIL pathway activity is significantly decreased in RMS compared with normal muscles in cohort 1 (wilcoxon-test, p-value < 0.0001) and cohort 2 (wilcoxon-test, p-value = 0.00155), while it is equivalently low in FN- and FPRMS (Fig. 1f and Extended Data Fig. 2e). This observation suggests similar blocking in apoptosis execution in both malignant entities. Since apoptotic pathways are predicted to be more activated in FNRMS, we then tested whether the expression profile of apoptotic effectors may have a clinical value in this subgroup. Using a cross-validation strategy, we generated an apoptotic gene metascore based on a two-genes signature with *BNIP3* (p-value < 0.05 in 186/250 iterations) and *FASLG* (p-value < 0.05 in 170/250 iterations) (see Methods; Extended Data Fig. 2f). We showed its association with the survival outcome of patients with FNRMS, with reproducible and significant effects in both independent cohorts 3 (HR = 2.7 [1.6-4.7]; p-value < 0.001) and 5 (HR = 1.7 [1.1-2.6]; p-value = 0.036) (Fig. 1g). We found that metascore-high patients, *i*.*e*. with a high expression of both BNIP3 and FASLG, have a significantly poorer outcome compared with metascore-low patients in FNRMS (log-rank tests; cohort 3, p-value < 0.001; cohort 5, p-value = 0.0028) (Fig. 1g and Extended Data Fig. 2g).

**Figure 1.**
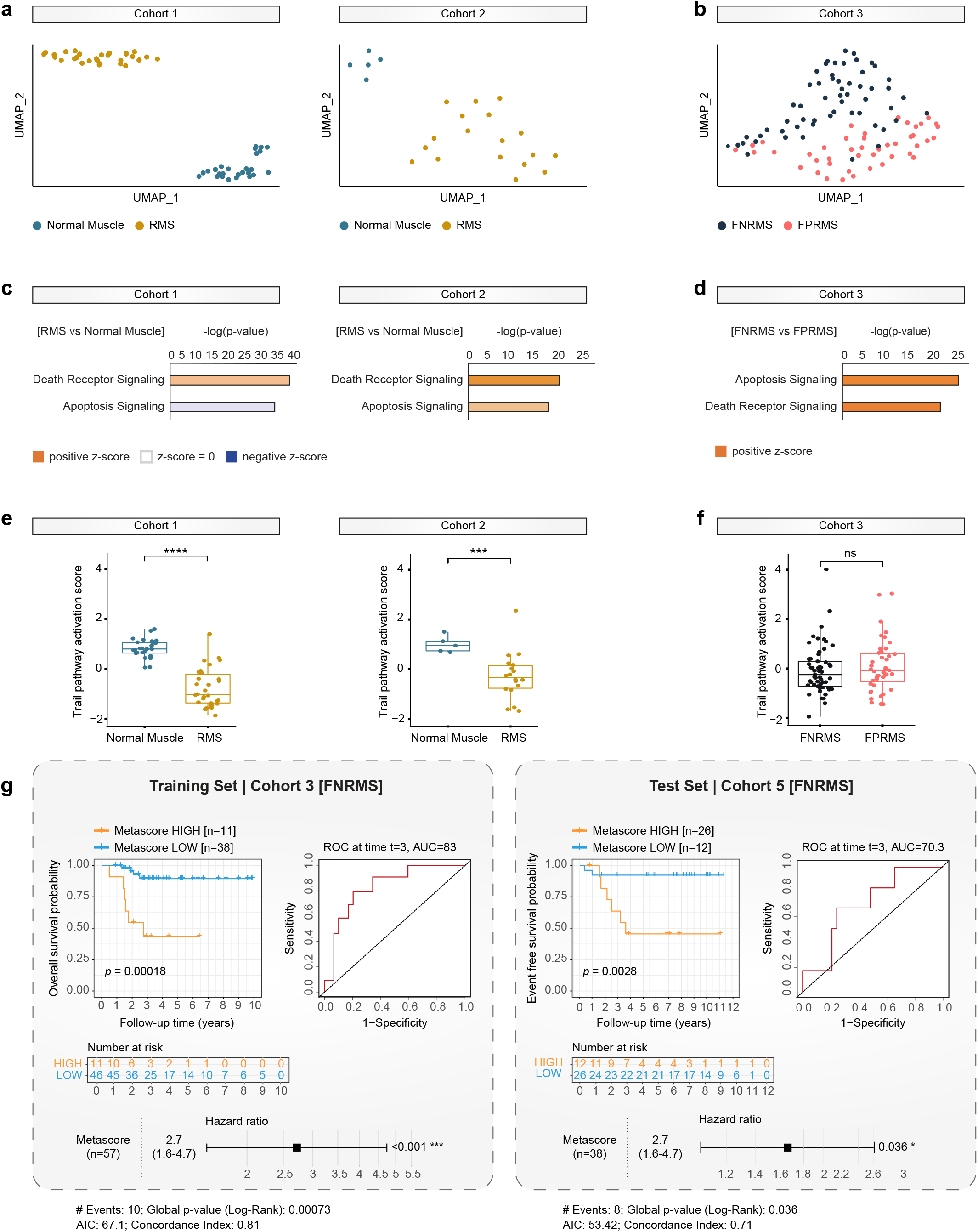
Apoptosis is committed and has a prognostic value in FNRMS. **a-b**. UMAP of RMS (yellow) and normal muscle (blue) samples (cohorts 1 and 2; **a**), or FNRMS (black) and FPRMS (pink) samples (cohort 3; **b**) based on the expression of apoptotic effectors (see Methods; Supplementary Table 1). **c-d**. Activation state of apoptotic cascades using Ingenuity Pathway Analysis in RMS versus normal muscle samples (cohorts 1 and 2; **c**) or FNRMS versus FPRMS samples (cohort 3; **d**). Significant difference corresponds to z-score > 0 and p-value < 0.0001. **e-f**. TRAIL apoptotic pathway activity inferred from a specific genes-response signature using PROGENy algorithm in RMS versus normal muscle samples (cohorts 1 and 2; **e**) or FNRMS versus FPRMS samples (cohort 3; **f**). Significant differences between groups are displayed on top for PROGENy analysis (wilcoxon signed-rank test; *** p-value ≤ 0.001; **** p-value ≤ 0.0001; ns, non significant). **g**. Survival analyses of the apoptotic metascore in FNRMS samples of the training (cohort 3) and test (cohort 5) sets. Kaplan-Meier curves were generated using its dichotomized form defined by a cross-validated optimal cutpoints procedure in a cohort-dependent manner. Differences of overall survival (cohort 3) and event free survival (cohort 5) probabilities between both groups were tested using log-rank tests and associated statistical probabilities are displayed on the graph. Number of patients at risk are indicated in the tables below the curves. Time-dependent receiver operating characteristic (ROC) curves and hazard ratios were generated using continuous apoptotic metascore. FNRMS: Fusion-Negative Rhabdomyosarcoma; FPRMS: Fusion-Positive Rhabdomyosarcoma; RMS: Rhabdomyosarcoma; UMAP: Uniform Manifold Approximation and Projection.

Overall, our transcriptomic profiling integration unveils a significant switch in the expression of apoptotic effectors between RMS and their non-tumoral muscle counterparts, which could be of prognostic interest to distinguish FNRMS high-risk patients. Causal inference of transcriptomic data predicts that apoptotic cascades are somehow committed in RMS, and notably in FNRMS, but fail to execute, underlying the necessity to pinpoint these blocking points from a therapeutic point-of-view.

### Overexpression of the anti-apoptotic *BIRC5* gene blocks FNRMS in a primed-for-death state

Given the predicted activation state of apoptosis in FNRMS, we sought to identify pathway’s blocking points. Differential gene expression analysis unveils that 72% of genes from a 86-gene apoptotic signature are differentially expressed (DE) between FNRMS and non-tumoral muscles (Fig. 2a and Supplementary Table 2). Interestingly, 47% of pro-apoptotic DE genes are expressed at significantly higher levels in tumors and 54% of the 13 anti-apoptotic DE genes are down-regulated in FNRMS compared with normal muscles. By plotting the master effectors of the extrinsic and intrinsic pathways that are DE on a signaling map, we observed that pro-apoptotic effectors overexpressed in tumors mostly belong to the upper part of the apoptotic cascade, consistently with our observation that apoptosis is committed in FNRMS (Fig. 2b). The expression pattern of genes encoding anti-apoptotic BCL-2 family proteins is contrasted, with a lower expression of *BCL2* in FNRMS compared with normal tissue (log2(FC) = −1.3; p < 0.0001), but a slight increase of *MCL-1* (log2(FC) = 0.6; p = 0.0015) and *BCL-XL* (log2(FC) = 0.6; p = 0.0016) expression, this gain being however not found in cohort 2 (Fig. 2a-b and Supplementary Table 2). Moreover, the pro-apoptotic *Bax* is significantly more expressed in FNRMS (log2(FC) = 2.9; p < 0.0001; Fig. 2a-b and Supplementary Table 2), suggesting that the pressure on apoptosis execution may not rely on the balance between pro- and anti-apoptotic BCL-2 family proteins. On the contrary, two major downstream pro-apoptotic effectors, the cytochrome C-encoding gene *CYCS* (log2(FC) = −0.8; p < 0.0001) and the caspase-independent endonuclease *ENDOG* (log2(FC) = −3.6; p < 0.0001) are less expressed in FNRMS (Fig. 2a-b and Supplementary Table 2), suggesting that the blockage of apoptosis execution is rather associated with a dysfunction of the downstream part. Accordingly, *BIRC5*, which encodes the IAP (Inhibitor of Apoptosis Protein) Survivin that inhibits the two executioners caspase-3 and −7, is significantly overexpressed in FNRMS (log2(FC) = 2.9; p < 0.0001) in two independent cohorts (Fig. 2a-c and Supplementary Table 2). This observation led us to consider that FNRMS may be primed-for-death. This notion is related to a state of cell dependency on an anti-apoptotic protein for survival: the corollary is that inhibition of this anti-apoptotic effector is sufficient to initiate cell death^32^. To define whether FNRMS may be dependent on *BIRC5* expression for survival, we performed a medium-scale drug screening focused on therapeutic compounds targeting apoptosis on 2D FNRMS cell lines (Fig. 2d). As expected, FNRMS cells are highly susceptible to the *BIRC5*-inhibitor YM155, with 10 nM of this compound being sufficient to drive massive cell death in all three tested 2D FNRMS cell lines (Fig. 2d and not shown), in agreement with the “primed-for-death” concept. In contrast, FNRMS cells are resistant to other IAP, BCL-2 and DNA repair inhibitors, confirming the blockage in apoptosis execution.

**Figure 2.**
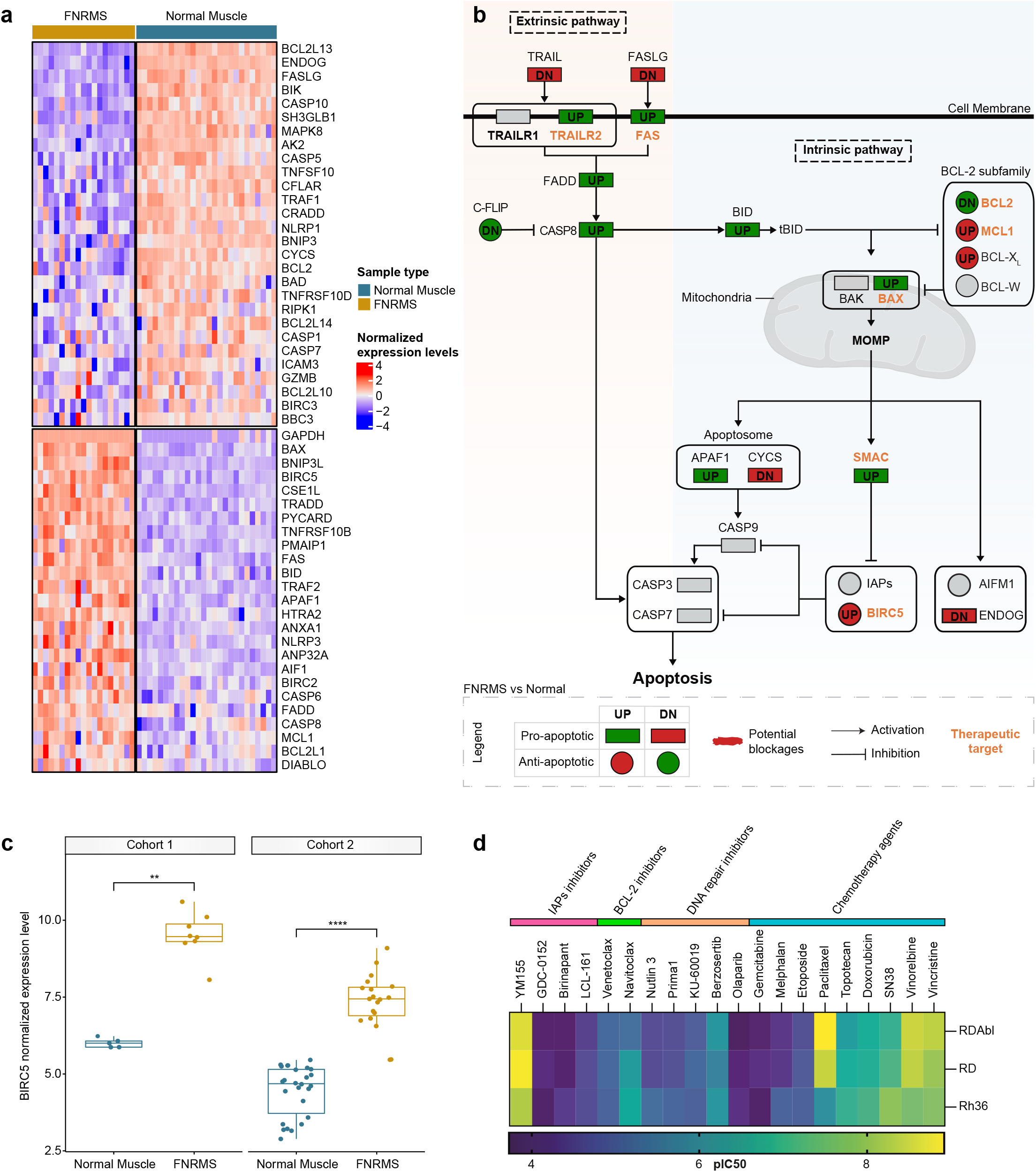
Overexpression of the anti-apoptotic *BIRC5* gene blocks FNRMS in a primed-for-death state. **a**. Heatmap representing the transcriptomic expression levels of apoptotic genes significantly differentially expressed (FDR < 0.05) between FNRMS (yellow) and normal muscle (blue) samples (cohort 1). Normalized and scaled gene expression levels are color-coded with a blue (low expression) to red (high expression) gradient. Samples in columns are clustered using Ward’s method on the inverse Spearman’s correlation coefficient matrix. **b**. Mapping of main pro-apoptotic (rectangle) and anti-apoptotic (circle) effectors significantly differentially expressed (FDR < 0.05; UP = FC > 1.5; DN [Down] = FC < 1.5) between FNRMS and normal muscle samples. Genes with altered expression compared with normal tissue are colored in red or green, when alterations are potentially associated with apoptotic blockage or not, respectively. Drug-target genes are indicated with an orange label. Abstract Inhibitors of Apoptosis (IAP)s comprise NAIP (BIRC1), BIRC2 (C-IAP1), BIRC3 (C-IAP2), XIAP (BIRC4). For a simplified overview, aliases are present including TRAIL (TNFSF10), TRAILR1 (TNFRSF10A, DR4), TRAILR2 (TNFRSF10B, DR5), C-FLIP (CFLAR), BCL-XL (BCL2L1), BCL-W (BCL2L2), SMAC (DIABLO). **c**. Expression levels of *BIRC5* between normal muscles and FNRMS samples (cohorts 1 and 2). Differences between groups were tested using wilcoxon signed-rank test with associated statistical probability displayed on top (** p-value ≤ 0.01; **** p-value ≤ 0.0001). **d**. Medium-scale drug screening of FNRMS cells (RDAbl, RD and Rh36) representing their sensitivity (pIC50) to a panel of 20 drugs, including IAPs, BCL-2, and DNA repair inhibitors, as well as chemotherapy agents, with a blue (low sensitivity) to yellow (high sensitivity) color-coded gradient. FC: Fold Change; FDR: False Discovery Rate; FNRMS: Fusion Negative Rhabdomyosarcoma.

Thus, integration of transcriptomic data and *in vitro* analysis of 2D cells sensitivity to apoptosis-targeting drugs support the notion that FNRMS can be considered as “primed-for-death”, relying on the expression of anti-apoptotic *BIRC5* for survival.

### FNRMS-derived organoids are new preclinical models that finely mimic tumor characteristics

Based on our results established on 2D FNRMS cell lines, *BIRC5* targeting could represent an appealing therapeutic strategy in FNRMS. However, lack of transposition of promising preclinical findings into efficient clinical treatments notably results from relying solely on such 2D cell lines, which do not faithfully reproduce the complexity of the tumors, notably in their heterogeneous and plastic components^33,34^. One of the ways to improve the predictive value of preclinical trials is to use models that recapitulate the biology of tumors and their behavior in the clinic as closely as possible. For this purpose, tumor-derived organoids (*i*.*e*. tumoroids) represent a considerable technological breakthrough. According to the initial definition^35^, these 3D cellular structures derived from tumor stem/progenitor-like cells, preserve the histological and molecular characteristics of their original tumor even after long-term expansion in culture, making them accurate and powerful tools for basic, translational and clinical research^35^.

Although new models were recently successfully derived from RMS patients’ biopsies or patient-derived xenografts^36,37^, such 3D-tumor-derived organoid models, characterized as reproducing the features of their original tumors and expandable over the long term to fit with the criteria of the organoid technology, have not been developed for FNRMS yet. By adjusting culture conditions according to the existing literature, but also by deciphering active signaling cascades that could support tumor cell growth thanks to transcriptome datasets (Extended Data Fig. 1), we have set up a culture medium (referred to as M3) and established a protocol that is sufficient to rapidly generate and expand 3D FNRMS-derived organoids (also designed thereafter as RMS-O or tumoroids) directly from fresh tumor specimens at relapse (100% efficiency, 6/6 samples; Fig. 3a and Supplementary Table 3). We have also used this protocol to derive primary 2D FNRMS models (100% efficiency on 7/7 samples, against 17% on 1/6 samples in classical DMEM-FBS 10% culture medium) (Fig. 3a and Supplementary Table 3). Keeping in mind that survival at relapse is less than 20% for patients with RMS, we decided to focus and characterize in depth these unprecedented 3D RMS-O models. All established RMS-O expand long-term (> 6 months) in culture, with a consistent split ratio of 1:2–1:3 every 10–15 days. These models are derived from tumors comprising diverse locations (head-neck, extremities, genito-urinary), histology (ERMS, pleomorphic RMS) and ages (pediatric and young adult patients) (Supplementary Table 3). RMS-O give rise to tumors when orthotopically xenografted (*tibialis anterior*) in mice, confirming their tumoral potential (Fig. 3b). RMS-O and their tumor-derived xenografts (RMS-XG) recapitulate accurately the histological features of their tumor-of-origin. For example, Patient 1 model (RMS1-O) preserves the patient’s tumor histology (RMS1-T), with cells of variable size, some with small nuclei and often reduced cytoplasm and some rhabdomyoblast cells, which are typical of RMS tumors, while the RMS-O derived from Patient 2 sample (RMS2-O) preserves the rather undifferentiated state of its tumor-of-origin (RMS2-T, Fig. 3b). Tumor morphological features are maintained in our 3D models even after long-term expansion in culture (over 6 months not shown). Besides histological organization, RMS-O also reproduce the expression pattern of key RMS diagnostic markers, such as Desmin and Myogenin, and the proliferation rates from their corresponding patient tissue (Fig. 3b).

**Figure 3.**
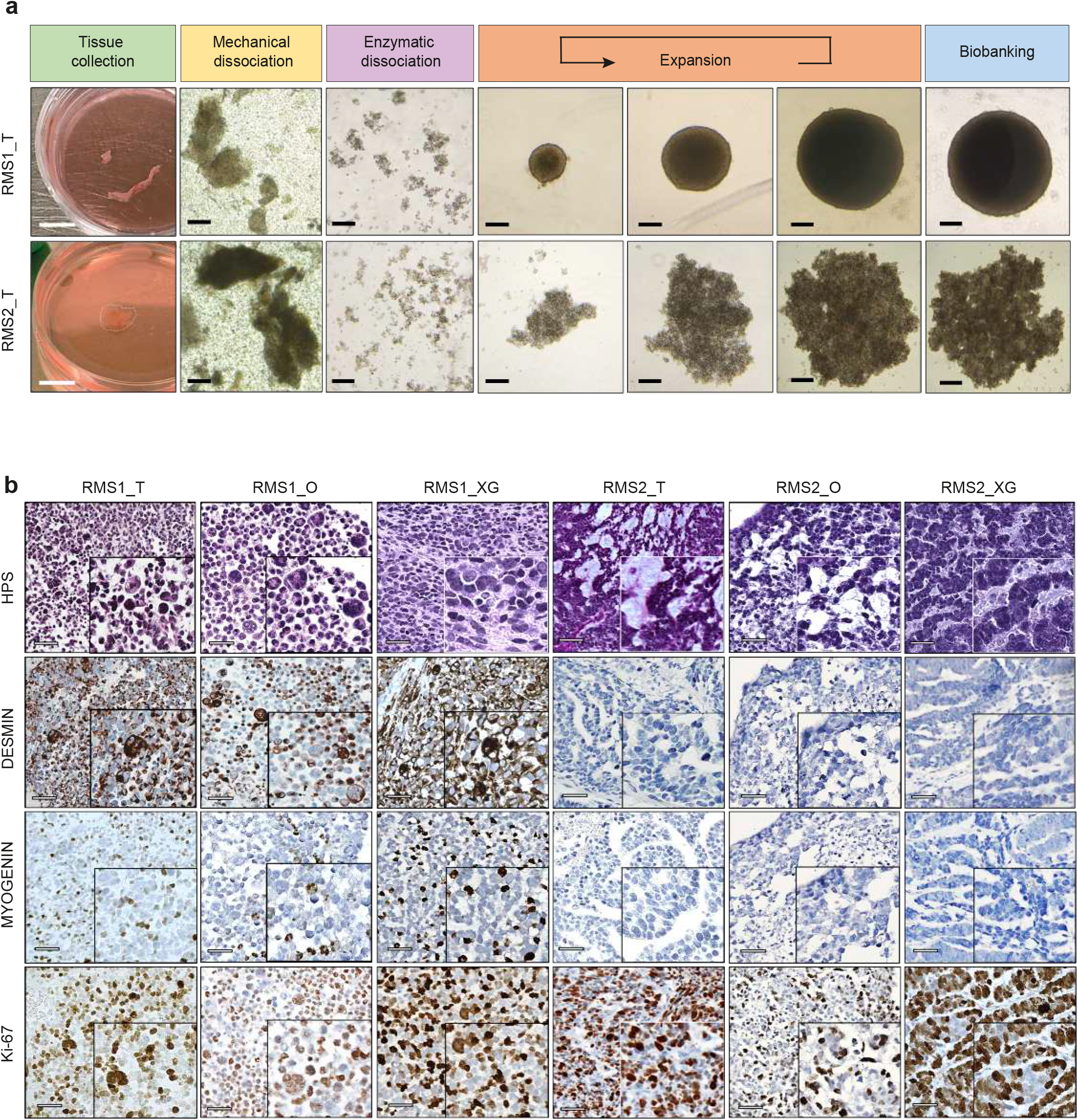
Design of a FNRMS-derived organoid model from fresh patients’ biopsies. **a**. Pipeline of organoids (RMS-O) derivation from fresh FNRMS tumors. Samples were obtained from patients undergoing biopsy/surgery (patients’ information detailed in Supplementary Table 3) and were processed as described in Methods. Six RMS-O have been established and expanded using this protocol. RMS1-O and RMS2-O derived from Patients 1 and 2, respectively, are shown. From day 3 to 15 post-seeding, FNRMS-derived organoids expand to 1,000 μm (RMS1-O) and 1,500 μm (RMS2-O) diameter. White scale bar: 1 cm. Black scale bar: 200 μm. **b**. Representative HPS and immunohistochemistry (IHC) characterization of RMS1-O and RMS2-O using clinical markers. RMS-O cultures and their matched-xenograft (RMS-XG) in mice were matched in blind by anatomopathologist experts to their tumor-of-origin (RMS-T). Expression of key clinical markers routinely used for RMS diagnosis such as Desmin, Myogenin and Ki-67 was evaluated by IHC. Scale bar: 50 μm. RMS1-T, Tumor from Patient 1; RMS2-T, Tumor from Patient 2; RMS1-O, Tumor-derived organoid from RMS1-T; RMS2-O, Tumor-derived organoid from RMS2-T; RMS1-XG, Xenograft from RMS1-O; RMS2-XG, Xenograft from RMS2-O. FNRMS, Fusion-Negative Rhabdomyosarcoma.

To further characterize FNRMS-derived organoid cultures and validate the adequation with their respective tumor-of-origin, we then compared their transcriptomic profiles using RNAseq analysis. An average of 100 million reads per sample were generated with 96% of the reads that mapped to the human genome of reference. While the first dimension explains most of the inter-patient heterogeneity, PCA analysis clearly shows that RMS-O models are grouped with their respective tumor-of-origin, both on Dim1 and Dim2 axes (Fig. 4a). On the contrary, corresponding 2D models cultured in standard DMEM-FBS 10% culture medium (DMEM_2D) or 3D models cultured in an incomplete culture medium (M2_3D) progressively derive from their initial tissue, simultaneously along Dim1 and Dim2 axes (Fig. 4a). Interestingly, we did not observe this phenomenon with 2D models grown in RMS-O optimized medium (M3_2D), even after long term growth and cryopreservation (Fig. 4a). Accordingly, Pearson’s correlation heatmap based on global gene expression profiles confirms that both RMS-O and their equivalent 2D models cultured in M3 medium (M3_2D) clusterize with their corresponding patient-derived tissues in an unsupervised analysis, while both DMEM_2D and M2_3D models are positioned in clusters independent of their tissue-of-origin (Fig. 4b). When examining more precisely RMS and differentiation markers, similar hierarchical clustering analysis highlights the high level of similarities between RMS-O and their corresponding tumor samples, even after cryopreservation, while unveiling notable differences with 2D cultures established from the same tumor sample, even if grown in M3 optimized medium (Fig. 4c). For example, for Patient 1 sample, the patterns of expression of both *GPC3* and *MYCN* were comparable in RMS-O and its tumor-of-origin, while being both downregulated in M3_2D culture. Similarly, the tumor from Patient 1 and its matched RMS-O, but not 2D cultures, both express markers of satellite cells and myoblasts such as *PAX7, MYOD1, TNNT2, MYL1*^38–42 43^ (Fig. 4c). These markers are barely expressed in both Patient 2 sample and its matched RMS-O, which reciprocally both present a reminiscent expression pattern of embryonic skeletal muscle development, as exemplified by the strong upregulation of *MEOX2*, a specific marker of early paraxial mesoderm^44^. FNRMS-derived organoid cultures then also accurately reproduce inter-patient heterogeneity and tumor differentiation status, and notably tumors’ spectrum of myogenesis markers’ expression. Moreover, functional gene set enrichment confirms that 2D cultures are characterized by unspecific processes notably related to cell adhesion regulation (Extended Data Fig. 3a). At the same time, FNRMS-derived organoids preserve an expected developmental and muscular identity (Extended Data Fig. 3a). Importantly, RMS-O also maintain more accurately, even after cryopreservation, the initial pattern of apoptotic gene expression compared to 2D models cultured in standard conditions (Extended Data Fig. 3b). Besides gene expression profile, RMS-O also faithfully retain the mutational landscape of the parental tumor, even after long term culture (>P20) and biobanking (Fig. 4d). Finally, as a proof-of-concept, we showed that RMS1-O reproduces the Vincristine resistance profile observed in Patient 1 (Fig. 4e-g). Although treatment is associated with a significant reduction in tumoroid growth and cell death (Fig. 4f-g), Vincristine treatment is insufficient to prevent the regrowth of tumor cells post treatment washout, which parallels the resistance observed in this patient (Fig. 4f-g).

**Figure 4.**
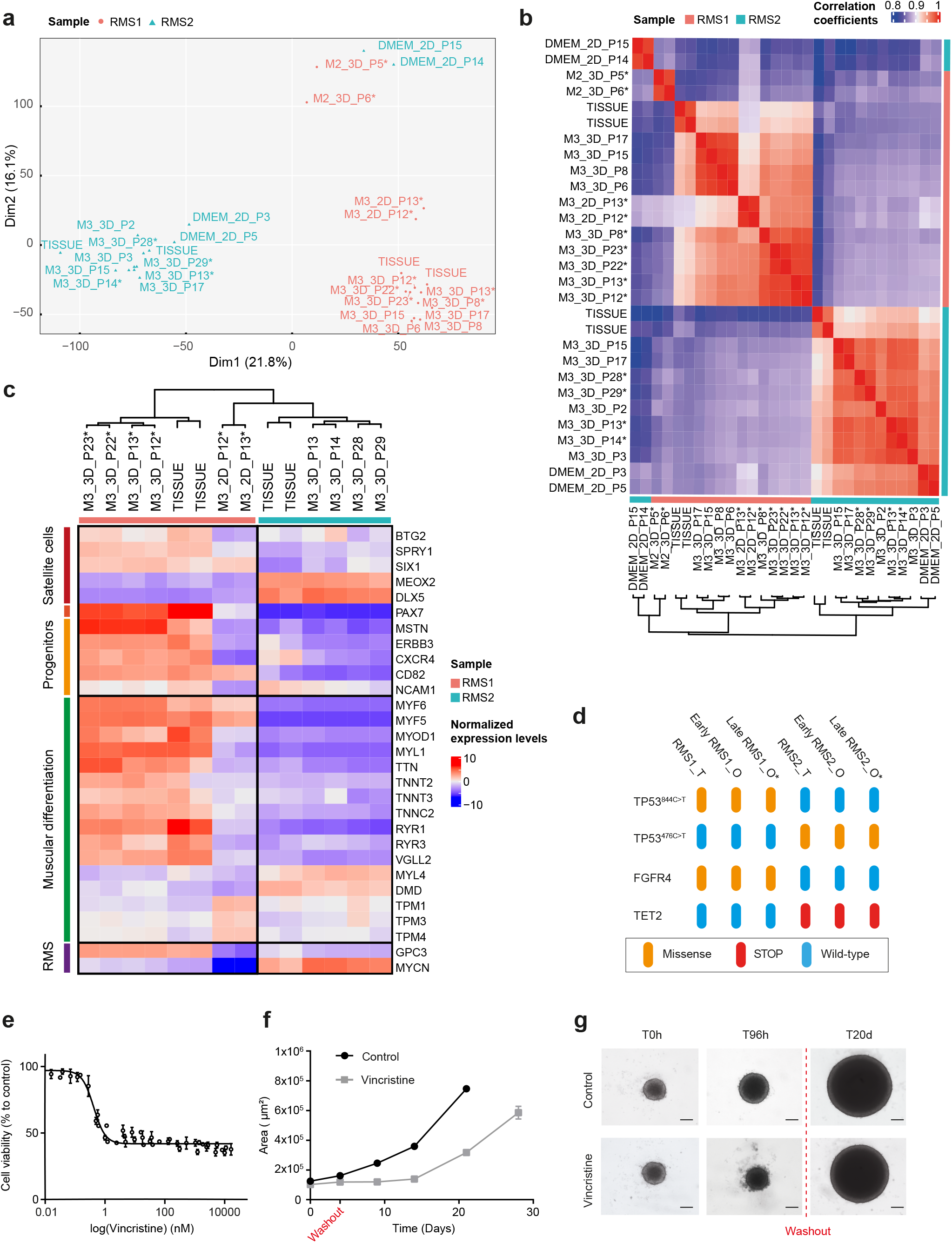
FNRMS-derived organoids are new preclinical models that finely mimic FNRMS characteristics. **a**. Principal Component Analysis (PCA) of RNA-seq data (normalized counts) plotted in 2D, using their projections onto the first two principal components (Dim1 and Dim2). Each data point represents one sample. Each sample is designed according to i) the medium in which it was derived, ii) its 2D or 3D structure, and iii) its passage at time of collection, and then labeled as follows: Culture Medium**_**Dimension**_**Passage. M3: optimized tumoroid medium; M2: incomplete medium; DMEM: Dulbecco’s Modified Eagle’s Medium. Patient 1-derived models and tissue (RMS1): pink dots; Patient 2-derived models and tissue (RMS2): blue dots. **b**. Pearson’s correlation heatmap based on global transcriptomic expression profile showing the clustering of RMS-O with their paired tissue-of-origin. Each sample is designed as above (see **a**). Color-coded annotation matches Patient 1-derived models and tissue as RMS1 (pink squares) and Patient 2-derived models and tissue as RMS2 (blue squares). **c**. Hierarchical clustering analysis based on the centered-normalized expression values of RMS tumor and differentiation markers highlights the high level of similarities between RMS-O and their corresponding tumoral samples. Top-left column indicates whether the indicated genes are markers of stem (progenitors/satellite cells) or committed muscle cells (muscle differentiation), or cancer features (RMS cancer). Each sample is designed as above (see **a**). Color-coded annotation matches Patient 1-derived models and tissue as RMS1 (pink squares) and Patient 2-derived models and tissue as RMS2 (blue squares). **d**. Preservation of tumor mutational profile in RMS-O. Sanger sequencing on gDNA of RMS-1 and −2 tissues and their corresponding tumoroids, respectively RMS1-O and RMS2-O, were performed after purification of PCR product surrounding mutations, based on genomics clinical reports. Early: RMS-O collected at early passages (<20). Late: RMS-O collected at late passages (>20). **e**. Vincristine dose-responses curves performed on FNRMS-organoid derived from Patient 1 (RMS1-O). Viability is expressed as a percentage of the value in untreated cells (CellTiter-Glo®). Means +/-std are represented (n=3). **f**. Growth curve of FNRMS-organoid derived from Patient 1 (RMS1-O) after treatment with Vincristine. RMS1-O were treated (grey line) or not (black line) with Vincristine during 4 days, before washout and follow-up of growth. Each point corresponds to mean +/-std of at least 3 RMS1-O areas. **g**. Representative brightfield images at 0 hour and 96 hours of treated (Vincristine) or control (CTL) RMS1-O. 1 nM; scale bar: 200 μm. FNRMS, Fusion-Negative Rhabdomyosarcoma; PCR, Polymerase Chain Reaction.

Therefore, these FNRMS-derived organoids are an unprecedented 3D-tool for studying RMS biology and response to treatments at relapse.

### The definition of relevant apoptotic-based therapeutic combinations relies on targeting FNRMS intra-tumor heterogeneity using tumoroids as models

Intra-tumoral heterogeneity is widely considered as a key driver of resistance to treatments^33,34^. To define whether our model preserves the recently described RMS tumor hierarchy reminiscent of human muscle development^37,45^, we generated droplet-based single-cell RNA sequencing (scRNA-seq) data on RMS-O using 10x Genomics’ technology. Overall, 14,371 tumoral cells (RMS1-0_P13 = 7608; RMS1-0_P14 = 6763) expressing a total of 33,538 transcripts and 15,175 detectable genes with more than 3 transcripts in at least one cell were quantified (Extended Data Fig. 4a). Biological replicates (different passages) were merged without batch correction as no difference was observed between samples (Extended Data Fig. 4b). We performed UMAP to visualize the unified transcriptomes of all cells^46^ and identified 6 clusters using an unsupervised Leiden algorithm^47^, each expressing a specific subset of biomarker genes (Fig. 5a; Supplementary Table 4). Supervised and unsupervised trajectory inference analyses on RMS-O scRNA-seq data unveils a myogenic differentiation sequence from quiescent satellite cell state (clusters 4-3) to myoblast-proliferative identity (clusters 5-2-6), accompanied by a gradual change in muscle differentiation markers (Fig. 5b-c and Extended Data Fig. 4c). Of note, clusters 5-2-6 define myoblasts at different cell cycle stages and were subsequently grouped into a single myoblast-proliferative cluster (Extended Data Fig. 4d). Interestingly, the intermediate cluster 1 fits with a human fetal skeletal muscle state already described in normal muscle^38^ (Extended Data Fig. 4e), which expresses canonical myogenic markers, albeit at slightly lower levels, but are mainly characterized by a gene signature suggesting a more mesenchymal-like nature^38^. We performed gene set enrichment analyses to confirm and further define cluster identities (Fig. 5d; Supplementary Table 5). Quiescent satellite cells are enriched in a hypoxic gene signature and in other genes involved in the regulation of ribosome and protein translation activity (Fig. 5d). On their side, cycling myoblasts are enriched in genes involved in quiescence exit, and in the previously reported pediatric cancer signature, which was generated by selecting DE genes between a panel of more than 70 xenografts from 8 types of childhood tumors (including RMS) and normal tissues^48^ (Fig. 5d).

**Figure 5.**
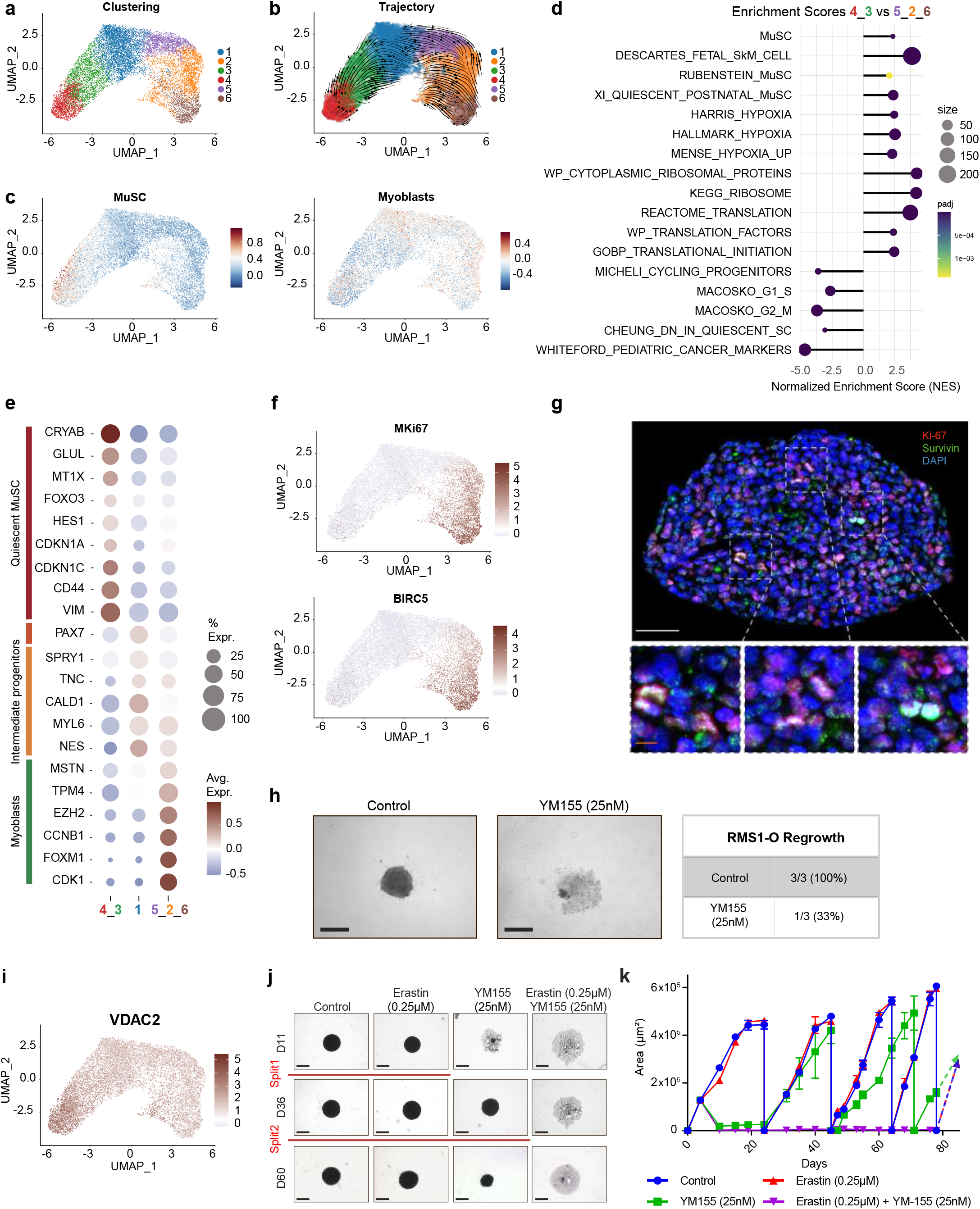
FNRMS-derived organoids preserve intra-tumor heterogeneity and can help improve therapeutic combinations targeting apoptosis. **a-b**. UMAP visualization of unified scRNA-seq data of tumoroids derived from Patient 1 (RMS1-O) samples (P13 and P14 passages) showing cluster identities (**a**), and unsupervised trajectory inference analysis using scVelo (**b**). **c**. Module scores of quiescent satellite cells (left panel, MuSC) and myoblasts (right panel) expression programs displayed on scRNA-seq UMAP of RMS1-O samples. **d**. Functional enrichment between quiescent satellite-like cells (clusters 4-3) and myoblast-proliferative cells (clusters 5-2-6) (see Methods). Dots are colored according to their adjusted statistical probabilities with a yellow (lower significance) to blue (higher significance) gradient and sized by the count number of genes matching the biological process. **e**. Dot plot representing gene expressions of specific myogenic differentiation markers between cluster groups. Dots are sized according to the percentage of cells in each cluster group that express the gene (transcript level > 0) and color-coded by average gene expression levels across cells. **f**. UMAP representation of *MKI67* (upper panel) and *BIRC5* (bottom panel) expressions in RMS1-O scRNA-seq data. **g**. Immunofluorescence showing heterogeneous expression of *BIRC5* (in red) among tumor cells on a tumoroid section (RMS1-O). Overlap between *BIRC5*-encoded protein, Survivin and Ki67 (in green) in proliferative cells is shown in enlarged boxes (bottom panel). Nuclei were counterstained with Hoechst 33342 (blue). White scale-Bar: 50 μm. Orange scale-Bar: 10 μm. **h**. BIRC-5 inhibitor YM155 shows transient efficiency on tumoroids. Left panel: representative images of FNRMS-derived organoids from Patient 1 (RMS1-O) treated or not with 25 nM of YM155 during two days. A halo of dead cells is visible around the residual cluster of tumor cells in the treated condition. Right panel: RMS1-O regrowth rate within 24 days after treatment washout, showing transient efficiency of YM155 in 33% of cases. Scale-Bar: 200 μm. **i**. UMAP representation of *VDAC2* expression in RMS1-O scRNA-seq data. **j-k**. Efficiency of Erastin/YM155 combination on RMS-O derived from Patient 1 (RMS1-O). RMS1-O were treated or not with YM155, Erastin or a combination of both for two days. Treatments were then stopped and regrowth of structures was evaluated within 80 days in each case. When regrowth of RMS1-O was observed, they were splitted two times to ensure that their renewal properties were preserved (**j**. Red lines, split ratio 1:2; **k**. vertical falls of growth curve lines to 0). **j**. Representative brightfield images of RMS1-O in the different conditions tested. **k**. Growth curve of RMS1-O after treatment washout in the different conditions tested. Each point corresponds to mean +/-std of at least 3 RMS1-O areas and lines are color-coded according to treatment: control (blue), Erastin (red), YM155 (green), combination of Erastin and YM155 (purple). UMAP, Uniform Manifold Approximation and Projection.

To go further, clusters were manually characterized according to their differential expression of key muscle marker genes described in the litterature^38–42^. As expected, clusters 4-3 express *CRYAB, GLUL* and *MTX1*, three genes robustly defining satellite cells in multiple recent scRNA-seq studies performed on human muscles^38,41,42^ (Fig. 5e). Cells in cluster 1 were characterized by markers of smooth muscle tissue like *CALD1* and *MYL6*^49^. Moreover, this cluster specifically express *PAX7*, suggesting a commitment towards myogenic differentiation and satellite cell activation^43^ (Fig. 5e). This differentiation ends at clusters 5-2-6, which express *EZH2*, a gene specifically induced in activated satellite cells during myogenesis^43^ and early myoblastic differentiation genes markers such as *MSTN* and *TPM4*^50^. Importantly, those tumor cells fail to express late markers of muscular differentiation (Supplementary Tables 4 and 5). Thus, proliferating myoblasts at the end of the myogenic continuum described in this RMS-O model appear as ‘halted’ in an early stage of differentiation. This confirms recent studies on RMS tumor hierarchy, showing that cells with different degrees of differentiation mirror the normal developmental stages, but fail to completely differentiate and remain stalled in actively cycling progenitor states^37,45^. We then asked if differences in apoptotic genes’ expression and associated drug sensitivity may exist in those different reminiscent developmental myogenic states. Interestingly, the proliferative halted-myoblasts population was specifically characterized by a high level of *BIRC5* expression (Fig. 5f). We validated this result by immunofluorescence on FNRMS-derived organoid and observed a colocalization between *BIRC5*-encoded protein Survivin, and Ki-67 in dividing cells (Fig. 5g). These data suggest that response to *BIRC5*-targeting drugs may be incomplete due to the heterogeneous expression of this apoptotic effector in RMS cells. To test this hypothesis, we evaluated the efficiency of *BIRC5*-inhibitor YM155 on this FNRMS-derived organoid model. RMS-O were exposed for 48h to a dose of YM155 sufficient to shut-off cellular ATP production as a marker of cell viability (Extended Data Fig. 4f). Treatment efficiency was evaluated after washout of the drug, by analyzing tumoroids regrowth. YM155 induces a massive wave of cell death and a major destruction of tumoroid structures at the treatment endpoint (Fig. 5h and Extended Data Fig. 4g). However, within 24 days post washout, 33% of tumoroids have grown back from leftover cells that were not eliminated by YM155 (Fig. 5h and Extended Data Fig. 4h). Based on our observation that *BIRC5* expression was restricted to the halted-myoblasts population, we looked for another target more strongly expressed in the quiescent satellite cell-like population. We identified the Voltage-Dependent Anion-selective Channel protein-2 encoding gene (*VDAC2*) as a putative candidate (Fig. 5i). VDAC2 forms a pore in the outer mitochondrial membrane and regulates its permeability to several molecules. This protein has been involved in the regulation of several processes and notably cell death *via* both ferroptosis and apoptosis modulation^51–53^. We then decided to evaluate the impact of a therapeutic combination, including YM155 and Erastin, a known inhibitor of VDAC2 activity^53,54^. Consistently with the pattern of expression of these genes in tumor cell clusters, this combined treatment is sufficient to induce a massive destruction of tumoroid structures in only 30h of treatment (Fig. 5j-k), at a dose of Erastin that is largely ineffective in monotherapy (Extended Data Fig. 4i). Most importantly, while Erastin or YM155-based monotherapies are largely insufficient in blocking RMS-O regrowth, their combination was on the contrary highly effective even after several weeks (Fig. 5j-k and Extended Data Fig. 4j).

Altogether, these data support that FNRMS-derived organoids are crucial tools to define effective therapeutic strategies in preclinical approaches, by pinpointing resistance resulting from the coexistence of different cell states.

## Discussion

Despite the implementation of many randomized trials from US and European groups, RMS remain a clinical challenge, due to the insufficient effectiveness of current treatments^16^. Therapy failure results from relapse due in part to intrinsic or acquired drug resistance. New tools are then necessary to improve the design of relevant drug combinations and to validate their efficiency in settings of robust preclinical trials. Here, we show that the use of organoid technology could make a major contribution to the definition and development of new therapeutic combinations, which are essential to improve the clinical outcome of patients with RMS.

Tumor-derived organoids have been almost exclusively derived from epithelial malignancies so far^55^. Here, we show that it is feasible to derive 3D-tumoroids from mesenchymal FNRMS tumors, which meet the definition of organoids since they accurately and precisely reproduce the histology and molecular characteristics of their tumor-of-origin. Although other protocols have been proposed^36,37^, our derivation pipeline is to date the only one that allows the accurate reproduction of FNRMS tumor features especially in their three-dimensional component, and the expansion of tumoroids over a long period of time. Preservation of this 3D structure is definitely a key issue to reproduce cell-cell contacts, as well as physical and mechanical constraints existing in malignancies, and thus to mimic tumor behavior^56^. Due to FPRMS tumor rarity, we were only able to test this derivation protocol on FNRMS to date. However, we believe that our pipeline is at least a solid basis to set up tumoroid models for other types of RMS, including FPRMS, although minor adjustments of the culture medium might be necessary. Moreover, the establishment of a collection of relapse FNRMS-derived organoids is highly relevant considering that i) this subgroup is the most prevalent one in children and adolescents, ii) a collection of such models, which we initiated here, is crucial to provide proxies of the high level of complexity resulting from the inter-patient heterogeneity existing in the FNRMS group and that iii) survival rate of patients at relapse is about 20%, pointing out the need for models to establish new efficient therapeutic approaches.

Indeed, one of the major challenges to improve RMS management is to define new therapeutic options that could be efficient for the largest fraction of FNRMS, despite the molecular diversity of this entity, and in particular for the high-risk forms, which still struggle to be clearly identified at the time of diagnosis^16^. Of note, we show here that FNRMS patients with a higher apoptotic metascore, established on the basis of apoptotic gene expression profiles, are significantly associated with a poorer prognosis. Several prognostic metascores have already been proposed to allow stratification of patients with FNRMS^57^, but none has been shown to be readily applicable in the clinic to date. Although tested on two independent cohorts, our mathematical model will have to be validated on larger groups of patients to define its clinical prognostic value compared to current criteria including tumor size, location or metastatic dissemination. It nevertheless highlights the fact that targeting apoptosis may still be relevant to improve the outcome of FNRMS patients.

Although a source of hope in the 2000s, targeting apoptotic pathways has so far shown limitations in preclinical testing and/or clinical trials, notably in RMS^26^. However, based on the integration of transcriptomic data, our results indicate that FNRMS can be considered as primed-for-death. Traditionally, programmed cell death was perceived as a binary black-or-white matter: cells are either alive or dead, depending on whether apoptotic pathways are turned on or off. This view has evolved to identify several cell states associated with intermediate activation of the apoptotic pathways. The notion of death priming has been used to define dependency of cancer cells on an anti-apoptotic protein for survival^58–60^. The underlying idea is that death pathways are committed but that their execution is prevented by overexpression of an anti-apoptotic effector, which until now was mainly described as a member of the BCL-2 family. Consequently, inhibition of this anti-apoptotic protein is sufficient to trigger apoptosis execution in these cells. Based on the exploitation of transcriptomic data and the use of apoptotic pathway activation signatures, our data suggest that upstream apoptotic pathways are indeed sustained in an activated state in FNRMS compared with their non-tumoral counterpart, *i*.*e*. skeletal muscle. This state results, in particular, from the overexpression of upstream pro-apoptotic proteins associated with the death receptor pathway, and from the loss of expression of major anti-apoptotic proteins such as Bcl-2. Although these data need to be confirmed at the protein level, considering the importance of both transcriptional and post-transcriptional mechanisms in the regulation of the expression of apoptotic effectors^61–64^, these findings argue for the identification of an anti-apoptotic protein whose inhibition would be sufficient to release the execution apoptosis in FNRMS cells.

Mapping of apoptotic pathways from bulk tumors’ transcriptomic data clearly suggested that targeting *BIRC5* could be this effective lever. The addiction state of FNRMS tumor cells to *BIRC5* overexpression was clearly confirmed *in vitro* by a medium-scale drug screening, focused on an apoptotic library of compounds. Indeed, 2D FNRMS cell lines appear highly sensitive to the *BIRC5*-inhibitor YM155. Targeting *BIRC5* has already been proposed as a putative therapeutic approach in several cancers including RMS, based on such *in vitro* assays^22,65^. New drugs targeting apoptosis are still being tested at early clinical study stage, more often combined with chemotherapy (NCT03236857; NCT04029688). However, to date, no combination including one of *BIRC5* inhibitors has shown clinical efficacy. Consistently, the use of our FNRMS-derived organoid models sheds new light on these contradictory data, and provides leads to re-explore the therapeutic potential of targeting *BIRC5* while explaining its limits and the potential origin of resistance observed in patients. Indeed, RMS-O models also preserve the intra-tumor heterogeneity of the original tumor. We were notably able to identify a developmental hierarchy within FNRMS comprising a quiescent satellite cell state, transitioning towards an activated mesenchymal-like state and halted-myoblasts, actively dividing and expressing early myogenic markers, thereby reproducing the incomplete muscle differentiation pattern described in RMS^37,45^. This reminiscent developmental program represents an interesting perspective to identify new therapeutic vulnerabilities that could be exploited to reduce disease recurrence. Very interestingly, *BIRC5* expression appears restricted to the myoblast proliferative clusters, consistently with the dual role of Survivin in both apoptosis inhibition and cell cycle promotion^66^. Then, the use of YM155, although associated with a robust decrease in FNRMS-derived organoid size and induction of cell death, only resulted in a transient effect in this model, with a resumption of growth observed upon discontinuation of treatment. Transposed to a patient context, *BIRC5* inhibition could then be only transiently sufficient, at clinically relevant doses, to trigger death of the proliferative fraction of tumor cells, but could be largely ineffective on the most quiescent stem ones, as previously observed with other therapies^6,37^. Bulk -omics characterization of tumors provides essential information on tumor biology and can be used to define the identity of tumor cell states. Notably, we observed an enrichment of the pediatric cancer signature defined by Whiteford and colleagues^48^ in RMS-O proliferative-myoblast clusters. Nevertheless, this observation also underlies the need to conduct single-cell omic characterization of tumors/models to unveil the full spectrum of tumor cell diversity, a prerequisite for defining efficient combined therapies^48^.

Because it finely reproduces the complexity and dynamics of intra-tumor heterogeneity, FNRMS-derived organoids then offer a seizable opportunity to reexplore the vulnerabilities of the tumor cell population that can be exploited therapeutically using relevant apoptotic-targeting drug combinations. The idea is not so much to define therapeutic synergies as to unveil complementarities based on targeting different tumor populations. We propose here YM155/Erastin as a putative therapeutic lead, but other promising strategies could be tested on our FNRMS models. In particular, PLK1 inhibition with volasertib has showed interesting efficacy *in vitro* on 2D cell lines and *in vivo*^67,68^– although tumor regrowth was observed in some cases–justifying its use in combination. Interestingly, *PLK1* expression is restricted to the G1/S proliferative cluster of FNRMS-derived organoids (data not shown), thus providing an explanation for the observed resistance and new potential combinations based on the simultaneous targeting of negative-*PLK1* populations. Besides apoptosis, this approach could be extended to other death pathways, especially considering the interconnection and plasticity of death cascades–some effectors being able to engage different signalings depending on the cellular context^7,8^–, and the targeting of different death effectors by several therapeutic compounds. Regarding this last point, we can not exclude here that Erastin exerts its death promoting effect solely *via* the inhibition of VDAC2, but also potentially *via* the induction of ferroptosis through depletion of GSH^53^.

Then, the use of FNRMS-derived organoid models could reconcile the promising data obtained *in vitro* and the failures observed in the clinic, in particular when targeting apoptotic pathways, to rapidly provide new effective therapeutic opportunities to prevent and anticipate resistance and relapse. The next challenge will be to establish a bio-collection of tumoroid models sufficient to mimic inter-patient heterogeneity and validate the relevance of these new approaches on a large fraction of patients.

## Methods

### Gene-Expression Analysis of available muscle/RMS datasets

Three microarray datasets were collected from public databases. Schäfer and Welle (cohort 1) log2-transformed data comprising 26 normal muscles and 30 RMS samples were downloaded from the R2 genomic platform (http://r2.amc.nl) using the gene reporter selection mode, *i*.*e*. HugoOnce algorithm that selects a single probeset to represent a gene. GSE28511 (cohort 2) quantile normalized data^69^ were downloaded from the GEO database (www.ncbi.nlm.nih.gov/geo/) and were then log2-transformed. After quality control, we removed the GSM706247 normal sample (tumor adjacent skeletal muscle cell) subject to high levels of tumor-in-normal contamination leading to a dataset of 5 normal muscles and 18 RMS samples. E-TABM-1202 (cohort 3)^13^ raw microarray data (.CEL files) with 101 RMS samples are accessible at the ArrayExpress platform (https://www.ebi.ac.uk/arrayexpress/) and were normalized using the Robust Multiarray Average (RMA) algorithm (oligo R library v.1.58.0). Last, Javed Khan and colleagues kindly shared Khan collection’s log2-transformed data (cohort 5) with 86 RMS samples^14^. Gene reporter selection was performed by selecting the probeset with the highest average expression levels across samples, except for the Schäfer and Welle dataset with default probeset assignment.

St. Jude RNA-seq data (cohort 4) of 60 RMS samples have been retrieved from St. Jude Cloud (https://www.stjude.cloud) and generated as described^70^. Briefly, read mapping was done using STAR (v.2.7.9a)^71^ on the hg38 human genome and gene-level counts were generated using HTSeq-count^72^ based on the Gencode v31 gene annotations^73^. We focused on transcripts with consistent annotations, *i*.*e*. protein-coding genes, and filtered those with less than 10 reads in overall samples. Gene expression data were normalized using a variance-stabilizing transformation procedure with vst function (DESeq2 R library v.1.34.0)^74^. To remove unwanted variability driven by technical and non-biological factors, we used the removeBatchEffect function implemented in the limma R library (v.3.50.3)^75^ and specified the “fusion status” as the variable to consider in the linear model.

### Apoptotic genes expression profiling and pathway activation scores

We selected manually curated genes, known to encode proteins involved in apoptosis and other forms of cell death mechanisms, from the Deathbase platform (http://deathbase.org/, downloaded on March 31, 2022). Only genes characterized in the *Homo sapiens* organism have been selected. Based on this list of 86 genes (Supplementary Table 1), we performed differential expression analyses using limma R library (v.3.50.3)^75^ for microarray data and DESeq2 R library (v.1.34.0)^74^ using Shrunken log2 fold changes (LFC) for RNA-seq data. We tested gene expression differences between (1) normal versus tumor samples in the cohort 1 (Schäfer and Welle) and cohort 2 (GSE28511); and (2) fusion-negative versus -positive tumor samples in the cohort 3 (E-TABM-1202), cohort 4 (St. Jude) and cohort 5 (Khan). Statistical probabilities were adjusted using the False Discovery Rate (FDR) method^76^. Only apoptotic genes with significant differences between the two conditions (FDR < 0.05) were then selected for visualization. Visualization plots were generated with the ComplexHeatmap R library (v.2.10.0) using ward.D2 clustering on the inverse Spearman’s correlation coefficient matrix to assess the distance between samples and genes. Single gene expression comparison between normal and tumor samples was performed using ggboxplot (ggpubr R library v.0.4.0) for visualization and rstatix (v.0.7.0) for statistical analysis using Wilcoxon signed-rank test. Ingenuity Pathway Analysis (IPA) was performed with QIAGEN IPA (v.01-20-04, https://digitalinsights.qiagen.com/IPA)^77^ to predict downstream effects on biological functions based on the expression log fold change ratio of apoptotic genes with significant differences between conditions (FDR < 0.05), *i*.*e*. normal versus tumoral or fusion-negative versus -positive samples. To infer TRAIL apoptotic pathway activity from gene expression data, we used progeny (R library v.1.16.0), a computational method built by analyzing large-scale transcriptomic changes from signaling perturbation experiments^78^.

### Establishment of a prognostic apoptotic metascore in patients with FNRMS

Only FNRMS patients with known survival time and status information were selected for analysis (cohorts 3 and 4). The cohort 3 (E-TABM-1202) was used as the training set and the cohort 4 (Khan) as the independent test set. For each apoptotic gene, univariate Cox proportional hazards models were performed to test the prognostic value of each gene. To limit optimism bias, the selection strategy was based on a leave-10-out cross-validation procedure with 250 iterations in the training set. Genes were ranked based on their statistical significance (p-value < 0.05) across iterations and those significantly associated with the overall survival probability of patients with FNRMS in at least 150 (60%) iterations were included in the multivariate Cox proportional hazards model. Proportional hazard hypothesis was checked using Schoenfeld residuals^79^ using cox.zph function (survival R library). In order to explore collinearity between predictor genes, associations were assessed with Pearson correlation coefficients using cor function (stats R library v.4.1.3) with method = “pearson”. For each sample, the apoptotic metascore was calculated as the sum of the predictor genes expression levels weighted by the regression coefficients of the training model, generated on the FNRMS samples of the cohort 1 (E-TABM-1202). For each cohort, an independent optimal risk cut point was identified in order to define two groups, high and low apoptotic metascore, among FNRMS. For each of the 250 iterations, a cut point of the metascore was identified using the surv_cutpoint function (survminer R library). This algorithm relies on the maxstat function (maxstat R library v.0.7-25) that performs a test of independence between a quantitative predictor X (here, the apoptotic metascore) and a censored response Y (here, the survival status) using maximally selected rank statistics. This defines which cutpoint μ in X determines two groups of observations regarding the response Y and measures the difference between the two groups as the absolute value of an appropriate standardized two-sample linear rank statistic of the responses. We retained as final threshold the median of overall cutpoints (n=250). The two groups defined by low and high apoptotic metascore have been studied in more detail from a discriminatory point of view. Kaplan-Meier survival curves were drawn using the ggsurvplot function (survminer R library). Survival curves in high and low metascore groups were compared using log-rank tests in the training and test sets. Dynamic receiver operating characteristic (ROC) curves were built using timeROC R library (v.0.4). All statistical analyses were performed in the R statistical environment (v.4.1.3) using survival (v.3.3-1; Therneau 2022), survivalROC (v.1.0.3; Heagerty 2013) and survminer (v.0.4.9; Kassambara 2021) libraries.

### Human specimens

Leftovers from RMS samples were obtained through biopsies/resections performed at the Pediatric Hematology and Oncology Institute (iHOPE, Lyon) or Hôpital Femme Mère Enfant (HFME, Lyon; AC2022-4937). The Biological Resource Centre (BRC) of the Centre Léon Bérard (n°BB-0033-00050) and the biological material collection and retention activity are declared to the Ministry of Research (DC-2008-99 and AC-2019-3426). Samples were used in the context of patient diagnosis. Non-used parts of the samples might be used for research if the patient is not opposed to it (information notice transmitted to each patient). This study was approved by the ethical review board of Centre Léon Bérard (N° 2020-02). This BRC quality is certified according to AFNOR NFS96900 (N° 2009/35884.2) and ISO 9001 (Certification N° 2013/56348.2). In brief, tumor pieces were put in a sterile saline solution (0.9%), while confirmed to be RMS by anatomopathologists. The study had all necessary regulatory approvals and informed consents are available for all patients. For each RMS sample, tissues were split into four parts and processed for histology, RNA and DNA isolation, or dissociated and processed for RMS models derivation.

### Derivation and culture of tumoroids and 2D lines

RMS tissues (∼5–125 mm^3^) were minced into small pieces, digested in a solution containing collagenase D (0.125 mg/mL Roche, cat. no. 1108866001) diluted in HBSS (Gibco, cat. no. 14025050) and washed using Advanced DMEM/F-12 medium (Gibco, cat. no. 12634010) supplemented with Hepes (1X, Gibco, cat. no. 15630106), GlutaMAX™ (1X, Gibco, cat. no. 35050038) and Penicillin-Streptomycin (1X, Gibco, cat. no. 15140122). After centrifugation, cultures were established in 96-well ULA plates (Corning, cat. no. 7007) or in 6-well plates (Corning, cat. no. 353046) either in DMEM or in an optimized M3 medium. Culture medium was changed twice a week, and RMS-organoids were split every 2 weeks on average. All cultures were tested every month for mycoplasma using the MycoAlert® Mycoplasma Detection Kit (Lonza, cat. no. LT07-318), in accordance with the manufacturer’s instructions. To prepare frozen vials, all organoid cultures were dissociated and resuspended in Recovery™ Cell Culture Freezing medium (Gibco, cat. no. 12648010).

### Xenograft

6-weeks-old male NSG-NOD SCID mice were obtained from Charles River animal facility. The mice were housed in sterilized filter-topped cages and maintained in the P-PAC pathogen-free animal facility (D 69 388 0202). For orthotopic grafts, 300 000 from RMS1-O and 500 000 from RMS2-O cells were prepared in 50% culture medium-50% Matrigel Low Growth Factor (Corning, cat. no. 356231) and were injected orthotopically into the *tibialis anterior* muscle of mice. Visible tumors developed in approximately 2-3 months (RMS1-O) and 3-4 weeks (RMS2-O). Mice were culled when the tumor reached the limit end-point (600 mm^3^). All experiments were performed in accordance with relevant guidelines validated by the local Animal Ethic Evaluation Committee (C2EA-15) and authorized by the French Ministry of Education and Research (Authorization APAFIS#28836).

### Histological analyses

FNRMS-derived organoids were fixed and processed as described before^80^. In brief, immunohistochemistry (IHC) was performed on an automated immunostainer (Ventana discoveryXT, Roche) using rabbit Omni map DAB kit. Organoids’ slides were stained with HPS (Hematoxylin Phloxine Saffron), or the following antibodies: anti-Desmin (1/50, Dako, cat. no. M0760), anti-Myogenin (1/100, Dako, cat. no. M3559), and anti-Ki67 (1/100, Dako, cat. no. M7240). Then, slides were incubated in relevant antibody-HRP conjugate for 1 hour at room temperature (RT) and finally revealed with 3,3′-diaminobenzidine (DAB) for 5 min, counterstained Gill’s-hematoxylin. Following IHC, slides were mounted using Pertex (Histolab, Ref# 00801-EX). Co-immunofluorescence (IF) was performed on Bond RX automated immunostainer (Leica biosystems) using OPAL detection kits (ref NEL871001KT, AKOYA bioscience). Primary antibodies specific to Survivin (1/400, Cell Signaling, cat. no. 2808S) and Ki-67 (1/100, Dako, cat. no. M7240) were applied 30 min at RT, as described previously^80^. Sequential immunofluorescence was performed using OPAL 520 (Survivin, green), OPAL 690 (Ki-67, red), and cells were counterstained with DAPI. Slides were then mounted in Prolong™ Gold Antifad Reagent (Invitrogen, Ref# P36930). Sections were scanned with panoramic scan II (3D Histech, Hungary) at 40× for IHC and using the Vectra POLARIS device (Akoya bioscience) for multiplexed IF.

### Molecular profiling of FNRMS-derived organoids

For RNA-seq library construction, 100 to 1000 ng of total RNAs from tissues/organoids were isolated using the Allprep DNA/RNA/miRNA universal kit (Qiagen, cat. no. 80224), RNeasy mini kit (Qiagen, cat. no. 74104) and Arcturus® PicoPure® RNA Isolation Kit (ThermoFisher Scientific, cat. no. KIT0204) following manufacturer’s instructions. Libraries were prepared with Illumina Stranded mRNA Prep (Illumina, cat. no. 20040534) following recommendations. Quality was further assessed using the TapeStation 4200 automated electrophoresis system (Agilent) with High Sensitivity D1000 ScreenTape (Agilent). All libraries were sequenced (2×75 bp) using NovaSeq 6000 (Illumina) according to the standard Illumina protocol.

Raw FASTQ files were then processed using the following steps. Quality control was performed using FASTQC (v.0.11.9)^81^, followed by trimming of adapter sequences with Cutadapt (v.3.4) using -a CTGTCTCTTATACACATCT and -A CTGTCTCTTATACACATCT parameters^82^. Reads were mapped using STAR (v.2.7.9)^71^ to the human reference genome assembly GRCh38.p13 with --seedSearchStartLmax 38 –outFilterMatchNminOverLread 0.66 --outReadsUnmapped Fastx --outSAMmultNmax -1 --outMultimapperOrder Random --outFilterScoreMinOverLread 0.66 --quantMode TranscriptomeSAM --outSAMstrandField intronMotif --twopassMode Basic --limitSjdbInsertNsj 1324910 parameters. Gene expression data was generated with HTseq-count (v.0.13.5)^72^ using --order pos --stranded reverse parameter and symbols were annotated with their respective Ensembl gene IDs using the package org.Hs.eg.db v3.14.0^83^ based on Gencode v37 (Ensembl v103)^73^.

Genomic DNA from both RMS tissues and models were extracted using the Allprep DNA/RNA/miRNA universal kit (Qiagen, cat. no. 80224) following manufacturer’s instructions. Polymerase Chain Reaction (PCR) were performed using the FIREPol® Master Mix Ready to Load (Solis biodyne, 04-12-00115) and a T100 thermal cycler (Biorad). Sequences of the primers used are the following ones: FGFR4_1648G>A (forward primer: 5’-TCTGACAAGGACCTGGCCGA-3’; reverse primer: 5’-CTCTCCTTCCCAGTCCTGGT-3’); TET2_220_C>T (forward primer: 5’-AACTTATGTCCCCAGTGTTG-3’; reverse primer: 5’-AGTCTGGCCAAAGAATGATC-3’); TP53_844C>T (forward primer: 5’-GGACCTGATTTCCTTACTCC-3’; reverse primer: 5’-GTGAATCTGAGGCATAACTG-3’); TP53_416C>T (forward primer: 5’-CTGTTCACTTGTGCCCTGAC-3’; reverse primer: 5’-CTGCTCACCATGGCTATCTG-3’). Amplified DNA were purified on a 1.5% agarose gel (Sigma-Aldrich, cat. no. 16500-500) and cleaned using the NucleoSpin© Gel and PCR clean-up kit (Macherey-Nagel, cat. no. 740609) following manufacturer’s instructions. Mutations were identified by Sanger sequencing performed by Eurofins genomics.

To assess the concordance of tissues with FNRMS-derived organoids, raw HTseq counts for all tissues and derived-models were loaded using DESeq2 R library^75^ with the “design” parameter combining sample conditions (tissue/culture, 2D/3D culture). Genes with low counts, *i*.*e*. less than 10 reads across samples, were then filtered. Gene expressions were normalized using the vst function of DESeq2 R library^74^ with parameter “blind=FALSE” and only protein coding genes were kept for further analysis. DESeq-normalized data were extracted using the DESeq function (DESeq2 R library). Principal Component Analysis (PCA) and Hierarchical Clustering on Principal Components (HCPC) were performed using FactoMineR (v.2.4)^84^ and factoextra (v1.0.7)^85^ R libraries. Heatmaps were generated using ComplexHeatmap R library (v2.10.0)^86^ with euclidean distance as the clustering method and color palettes of RcolorBrewer R library (v.1.1-3). All analyses were performed in a R statistical environment (v.4.1.2) using DESeq2 (v1.34.0) library.

### Single-cell RNA sequencing analysis of FNRMS-derived organoids

For single-cell suspension preparation, FNRMS-derived organoids were dissociated using TrypLE Express Enzyme (ThermoFisher Scientific, cat. no. 12605010) preheated to 37 °c for 3 min. Cells were then filtered through a 30-μm strainer (Miltenyi Biotec, cat. no. 130-098-458), centrifuged at 500g for 5 min, resuspended in complete culture medium and sorted using a FACSAria (BD Biosciences). Cells were centrifuged again at 500g for 8 min and resuspended in PBS (Gibco, cat. no. 14190-094) with 0.04% BSA (Sigma-Aldrich, cat. no. A7030) for a final cell concentration of 1,000 cells/μL. Approximately 10 μL of isolated cells were loaded on a 10X Genomic chip and run on the Chromium Controller system (10X Genomics) to target 10,000 cells per sample.

Gene expression data was generated with the Chromium Single Cell 5’ v3.1 assay (10X Genomics) and sequenced on the NovaSeq 6000 platform (S1 flow cell, Illumina). To generate gene-barcode count matrices, raw sequencing reads were processed using mkfastq and count (Cell Ranger v.3.1.0, 10x Genomics)^87^. The raw base call (BCL) files were demultiplexed into FASTQ files and aligned to the hg38 human genome as reference. Overall, 23993 cells (RMS2_P13, n=11627; RMS2_P14, n=12366) passed the quality control criteria. Each single cell dataset was imported using Read10X function and converted into a Seurat object with CreateSeuratObject function with at least min.features = 200 and min.cells = 3. To retain only high-quality cells, we applied a joint filtration based on unique molecular identifier (UMI), number of detected genes and number of mitochondrial counts criteria^88^. For each sample, independently, we retained cells within a three median absolute deviation (MAD) around the population median for these metrics, combined with absolute quality thresholds. We considered low-quality cells as cells with (1) low (nGene < 200 genes) and high (nGene > 3MAD) number of detected genes; (2) high mitochondrial gene content (mitoRatio > 3MAD); and (3) cells with relatively high library sizes (nUMI > 4,500). We predicted doublets/multiplets, *i*.*e*. multiple cells captured within the same droplet or reaction volume, using the scDblFinder R library (v.1.10.0)^89^ but kept this variable as indicative. The single-cell datasets were then merged and normalized using methods adapted from Scran pipeline (scran R library v.1.24.0) comprising quickCluster, computeSumFactors with min.mean = 0.1 and logNormCounts steps^90^. The highly variable genes (HGVs) were detected using three algorithms including scran, Seurat and a rank custom strategy. The scran method comprises: (1) a modelGeneVar function that models the variance of the log-expression profiles for each gene; (2) a metadata function to fit the mean-variance trend; and (3) a getTopHVGs function to extract the top features. The Seurat V3 algorithm was implemented in the highly_variable_genes function (scanpy python library v.1.8.2)^91^ and consists of ranking genes according to a normalized variance procedure. The custom strategy (1) ranks genes according to their expression levels for each cell; (2) measures the standard deviation of rank for each gene across overall cells; and (3) sort genes based on their ranked expression levels; and (4) select the most variable ones. For each strategy, we selected the top 2,000 most variable genes and retained a list of 1,158 genes that were detected in at least two of the three methods. Variable features included the top 484 of these most variable genes and 245 genes known to be biologically relevant in the process of myogenic differentiation^38,41,42,92,93^ and were used for principal component analysis (PCA) using RunPCA function. We kept the first 9 principal components (PCs) for analysis based on the ElbowPlot method that allows a visualization of the standard deviation of each PC. The most contributing dimensions were then chosen based on two metrics: (1) the percent of variation associated with each PC (cumulative percent of variation > 90% and percent of variation < 5%); and (2) the percent change in variation between consecutive PCs (< 0.1%). Clusters were identified with the FindClusters function using a resolution set to 0.3 and the Leiden algorithm. Briefly, this strategy comprises local moving of nodes, refinement of the partition and aggregation of the network based on the refined partition, as previously described^47^. Cluster identities of the cells were then mapped on a UMAP using the RunUMAP function. Specific marker genes for clusters were identified using the FindAllMarkers function with only.pos = TRUE, min.pct = 0.25 and test.use = “MAST”. Trajectory inference analyses was performed using slingshot R library (v.2.4.0)^94^ with start.clus = 4 and stretch = 0 for a supervised strategy and scVelo python library (v.0.2.4) for an unsupervised one based on RNA velocity data generated by loom python library (v.3.0.6). For each cluster, we also identified both positive and negative cluster marker genes using FindAllMarkers function with min.pct = 0.25 and test.use = “MAST”. We then ranked these genes based on their fold change ratio and performed functional enrichment analysis. HALLMARK (H), Gene Ontology (subcategory: Biological Processes), curated (C2) and cell type signature (C8) gene sets, downloaded from MSigDB (http://www.gsea-msigdb.org/)^95^, and custom lists based on literature review (Supplementary Table 5) were selected for functional analyses. Overall, 14,818 gene sets were tested for Gene Set Enrichment Analysis (GSEA) using fgsea R library (v.1.22.0). Statistical probabilities were adjusted based on the number of tested biological processes using the FDR method^76^. Only custom, hypoxic, ribosomal and translational biological processes with a significant enrichment (FDR < 0.01) were retained for Fig. 5d. Analyses were performed in a R statistical environment (v.4.1.3) using Seurat R library (v.4.1.1)^96^ and python environment (v.3.9.10).

### Analysis of drugs’ impact on FNRMS-derived organoids

For IC50 determination, tumoroids were dissociated and plated at 5000 cells/well in 96-well ULA plates (Corning, cat. no. 4515). RMSO were allowed to form during 72 hours, and then treated with serial dilutions of YM155 (Selleckchem, cat. no. S1130), Erastin (Selleckchem, cat. no. S7242) or Vincristine (Teva, collected at the pharmacy of Centre Léon Bérard). Impact of treatments on intracellular ATP content was measured using the CellTiter-Glo® 3D Cell Viability Assay (Promega, cat. no. G9681) at 48 hours (Erastin/YM155) or 96 hours (Vincristine). Relative luminescence units (RLU) of each well were normalized to the mean RLU from the DMSO negative control wells as 100% viability. Gambogic acid (10 μM, Cayman Chemical, cat. no. 14761) was used as a positive control. All acquisitions of luminescence were performed on a Spark® microplate reader (Tecan) with a 400 ms exposition and auto-attenuation. Three technical replicates per condition were performed for each experiment. IC50 curves were drawn using Prism 7.04 (GraphPad).

For washout experiments, tumoroids were seeded at 5000 cells/well in 96-well ULA plates (Corning, cat. no. 7007). After 72 hours, RMSO were treated either with DMSO as negative control, or 0.25 μM Erastin, or 25 nM YM155, or a combination of both compounds. Two days after, RMS-O were then collected, washed twice in fresh medium and put back in new wells with complete culture medium. Culture medium was renewed once a week. When reaching the growth plateau (around 4 to 6×10^5^ μm^2^ in area), FNRMS-derived organoids were split and reseeded at 5,000 cells/well.

For IF staining of dead and viable cells, the LIVE/DEAD™ Viability/Cytotoxicity Kit (ThermoFisher Scientific, cat. no. L3224) was used directly on treated RMS-O following manufacturer’s instructions. RMS-O imaging was performed using the EVOS™ M7000 Imaging System (Invitrogen).

### Drug screening on 2D cell lines

Briefly, 2,000 living cells from RDAbl and 4000 from RD or Rh36 FNRMS cell lines were seeded per well in 384-well plates (Corning, cat. no. 3830) and incubated in the presence of a selection of 20 drugs in a humidified environment at 37°C and 5% CO2. Cells were grown in DMEM medium supplemented with 10% fetal bovine serum, 1% penicillin-streptomycin, 1% Glutamax and 1% non-essential amino acids. Drugs were distributed with the Echo 550 liquid dispenser (Labcyte) at 6 different concentrations covering 3 logs (*i*.*e*., 1 nM to 1 μM; 10 nM to 10 μM; 100 nM to 100 μM) in constant DMSO. Cell viability was measured using CellTiter-Glo® 2.0 Cell Viability Assay (Promega, cat. no. G9243) after 72 hours of drug incubation and luminescence was read using a Pherastar® plate reader (BMG Labtech). Data were normalized to negative control wells (DMSO only). IC50, defined as half maximal inhibitory concentration values, were deduced from dose-response curves obtained using Prism 9.3.1 (GraphPad).

## Data availability

Raw bulk and single-cell RNA sequencing data will be available *via* GEO. Accession numbers are pending. Supplemental Tables 4 and 5 are available upon request.

## Code availability

All data analyses’ codes will be made available without restriction upon request.

## Contributions

C.S, P.H. L.L., A.T., J-Y.B., M.C. and L.B. conceived and designed the experiments and analyses. M.C. and L.B. analysed the data with contributions from C.S, P.H. L.L., A.T., C.D., J-Y.B. and T.D. Bioinformatic analyses were performed by C.S. and T.D., with the help of L.T., A.B., P.R, C.D., N.M., V.M., P.R., M.C. and L.B.. Tumor samples processing and characterization was performed by L.L., C.D., and L.B., with contributions from I.R., N.M., C.C, C.P., F.D, C.B. and N.C. Metascore design and statistical analysis were performed by C.S., M.C. and D.M.B.. M.L.G., K.M., and E.P. performed drug screening on 2D cell lines, while P.H. and A.T. did drug experiments on tumoroids. M.C.B and A.D. provided detailed technical advice. Sequencing was performed on the CRCL platform with the help of C.D. and V.A. Histological and immunostaining were performed with the help of the anatomopathological platform of CRCL with the help of S.L. and N.G. M.C. and L.B. wrote the manuscript with contributions from C.S, P.H. L.L., A.T., C.D. and J-Y.B. M.C. and L.B. supervised the work. All authors read and approved the manuscript.

## Acknowledgements

We thank the patients and their families who consent to participate in this study, as well as parents’ charities which support this work (“Enfants, Cancers, Santé”, “Imagine for Margo”, “L’Etoile de Martin”, “Le sourire de Lucie”, “Aidons Marina”, “Un Horizon d’Espoir”, “Nos P’tites Etoiles”, and “César Gibaud, une enfance sans cancer”). We also thank clinical teams from IHOPe and HFME for their support and contributions, as well as all the facilities from the CLB and CRCL. We are grateful to Séverine Tabone-Eglinger, Loïc Sebileau and Anne-Sophie Bonne for their help with the management of regulatory procedures. We also thank Dr Xavier Morelli and Carine Derviaux of the HiTS platform of CRCM for their help with the medium-throughput drug screening. This research was supported by the Foundation ARCECI innovation award (USA) awarded to L.B., the ANR JCJC program, the DevWeCan2 LabEx, the Convergence Institute Plascan and the Ligue Nationale Contre le Cancer. C.S. and A.T. received financial support respectively from the Fondation de France and the Ligue Nationale Contre le Cancer.

