## Supplemental Figures for "Fusion-negative Rhabdomyosarcoma 3D-organoids as an innovative model to predict resistance to cell death inducers"

#### Extended Data Figures Legends

**Extended Data Figure 1.** Clinical description of the five cohorts used for transcriptomic analyses. Samples proportions depending on the clinical categories are represented: gender, age group (in years), histology, fusion status, clinical status in the RMS/normal muscle (cohorts 1 and 2) and RMS only datasets (cohorts 3, 4 and 5). ARMS: alveolar rhabdomyosarcoma; ERMS: embryonal rhabdomyosarcoma; FDR: False Discovery Rate; FNRMS: Fusion-Negative Rhabdomyosarcoma; FPRMS: Fusion-Positive Rhabdomyosarcoma; RMS, Rhabdomyosarcoma.

**Extended Data Figure 2. a.** PCA of apoptotic effectors' transcriptional levels (see Methods; Supplementary Table 1) from cohorts 1 and 2, plotted in 2D, using their projections onto the first two principal components (Dim1 and Dim2). Each dot represents one sample with normal muscles in blue and RMS in yellow. The first principal component (Dim1) is strongly correlated with the status (normal/tumoral) of samples. **b.** UMAP of FNRMS (black) and FPRMS (pink) samples (cohort 4) based on the expression of apoptotic effectors (see Methods; Supplementary Table 1). **c.** PCA analysis of apoptotic effectors' transcriptional levels (see Methods; Supplementary Table 1) from cohorts 3 and 4, plotted in 2D, using their projections onto the first two principal components (Dim1 and Dim2). Each dot represents one sample with FNRMS in black and FPRMS in pink. The second principal component (Dim2) is correlated with the status (FP/FN) of samples. **d.** Activation state of apoptotic cascades using Ingenuity Pathway Analysis Software in FNRMS versus FPRMS (cohort 4). Significant difference corresponds to z-score > 0 and p-value < 0.0001. **e.** TRAIL apoptotic pathway activity inferred from a specific genes-response signature using PROGENy algorithm in FNRMS versus FPRMS (cohort 4). Significant differences between groups are displayed on top of PROGENy analysis (wilcoxon signed-rank test; ns, non significant). **f.** Apoptotic metascore between alive (black) and dead (grey) patients with FNRMS in the cohorts 3 and 5. Differences between groups were tested using wilcoxon signed-rank test and associated statistical probability are displayed on top (* p-value ≤ 0.05; *** p-value ≤ 0.001). **g.** Workflow for construction and evaluation of an apoptotic gene signature and metascore in a training set (cohort 3) and independent test set (cohort 5). FNRMS: Fusion-Negative Rhabdomyosarcoma; FPRMS: Fusion-Positive Rhabdomyosarcoma; PCA: Principal Component Analysis; UMAP: Uniform Manifold Approximation and Projection.

**Extended Data Figure 3. a.** Functional enrichment of DE genes between 2D cell lines and 3D-RMS-derived organoid from Patient 1. Left scatterplots represent the top 6 enriched Gene Ontology (GO) pathways in 2D models, right scatter plots corresponding, reciprocally, to the top 6 enriched Gene Ontology (GO) pathways in 3D FNRMS-derived organoids. Dots are colored according to their adjusted statistical probabilities with a blue (lower significance) to yellow (higher significance) gradient and sized by the count number of genes matching the biological process. **b.** Hierarchical clustering analysis based on the centered-normalized expression value of apoptotic genes highlights the high level of similarities between tumoroids and their corresponding tumor samples. Top-left column indicates whether the indicated genes are markers of stem (progenitors/satellite cells) or committed muscle cells (muscle differentiation), or cancer hallmarks (RMS Cancer). Each sample is designed as follows, according to i) the medium in which it was derived, ii) its 2D or 3D structure, and iii) its passage at time of collection: Culture Medium_Dimension_Passage. M3: optimized tumoroid medium; M2: incomplete medium; DMEM: Dulbecco's Modified Eagle's Medium. Patient 1-derived models and tissue (RMS1): pink square; Patient 2-derived models and tissue (RMS2): blue square.

**Extended Data Figure 4. a-e.** UMAP visualization of unified scRNA-seq data of FNRMS-organoid derived from Patient 1 (RMS1-O) samples. (**a**) Quality control metrics (nGene, number of detected genes; nUMI, number of transcripts; and mitoRatio, percentage of mitochondrial transcripts). (**b**) Biological replicates (RMS1-O samples at P13 and P14 passages). (**c**) Unsupervised trajectory inference analysis using slingshot. (**d**) Module scores of cycling progenitors, G1_S and G2_M cell cycle phases, from left to right. (**e**) Module scores of fetal skeletal muscle and mesenchymal cells. **f.** IC50 curve representing RMS1-O sensitivity to YM155. Viability is expressed as a percentage of the untreated condition (CellTiter-Glo) in function of log2(drug concentration). Red dot indicates the concentration that was used in further experiments. **g-h.** Immunofluorescence staining of dead (red) and viable (green) cells in RMS1-O treated or not with YM155. Brightfield images are shown just after the treatment was stopped **(g)** or after regrowth **(h)**. Fluorescent ethidium homodimer-1 (orange) and calcein AM (green) respectively allow to distinguish dead and live cells in treated (bottom panel) versus control (upper panel) RMS1-O. Scale Bar: 200µm. RMS1-0: Rhabdomyosarcoma-organoid derived from Patient 1. **i.** IC50 curve representing FNRMS-derived organoid from Patient 1 (RMS1-O) sensitivity to Erastin. Viability is expressed as a percentage to the untreated condition (CellTiter-Glo) in function of log2(drug concentration). Red dot indicates the concentration that was used in further experiments. **j.** RMS1-O regrowth rate within 80 days after treatment washout, showing improved efficacy of the combination Erastin/YM155. UMAP: Uniform Manifold Approximation and Projection.

#### Supplementary Tables Legends

**Supplementary table 1.** Apoptotic genes list from the Deathbase database.

**Supplementary table 2.** Differential gene expression analysis of apoptotic genes in cohorts 1 to 5.

**Supplementary table 3.** Evaluation of the optimal media for 3D-RMS-O and 2D cultures of FNRMS and patients' clinical characteristics. M3 culture condition was defined as optimal based on its suitability to sustain continuous RMS-O growth. For 2D cultures,

RMS-O like and DMEM culture conditions were defined as optimal or non-optimal based on their suitability to sustain continuous proliferation of cells and to support cell growth after a freezing/thawing cycle. NA, Not Applicable; ND, Not Documented.

**Supplementary table 4.** Cluster biomarkers of the scRNA-seq analysis of RMS1-O model with genes being tested for differential expression between one cluster versus all the others.

**Supplementary table 5.** Functional enrichment of RMS1-O scRNA-seq data. (**a**) Custom gene-sets used for characterizing the myogenic differentiation programs of RMS1-O scRNA-seq data. (**b**) Functional enrichment of RMS1-O scRNA-seq data between quiescent satellite cell-like cells (clusters 4-3) and myoblast-proliferative (clusters 5-2-6) populations using Gene Set Enrichment Analyses (GSEA) based on differential expression genes.

### Extended Data Figure 1

**a**

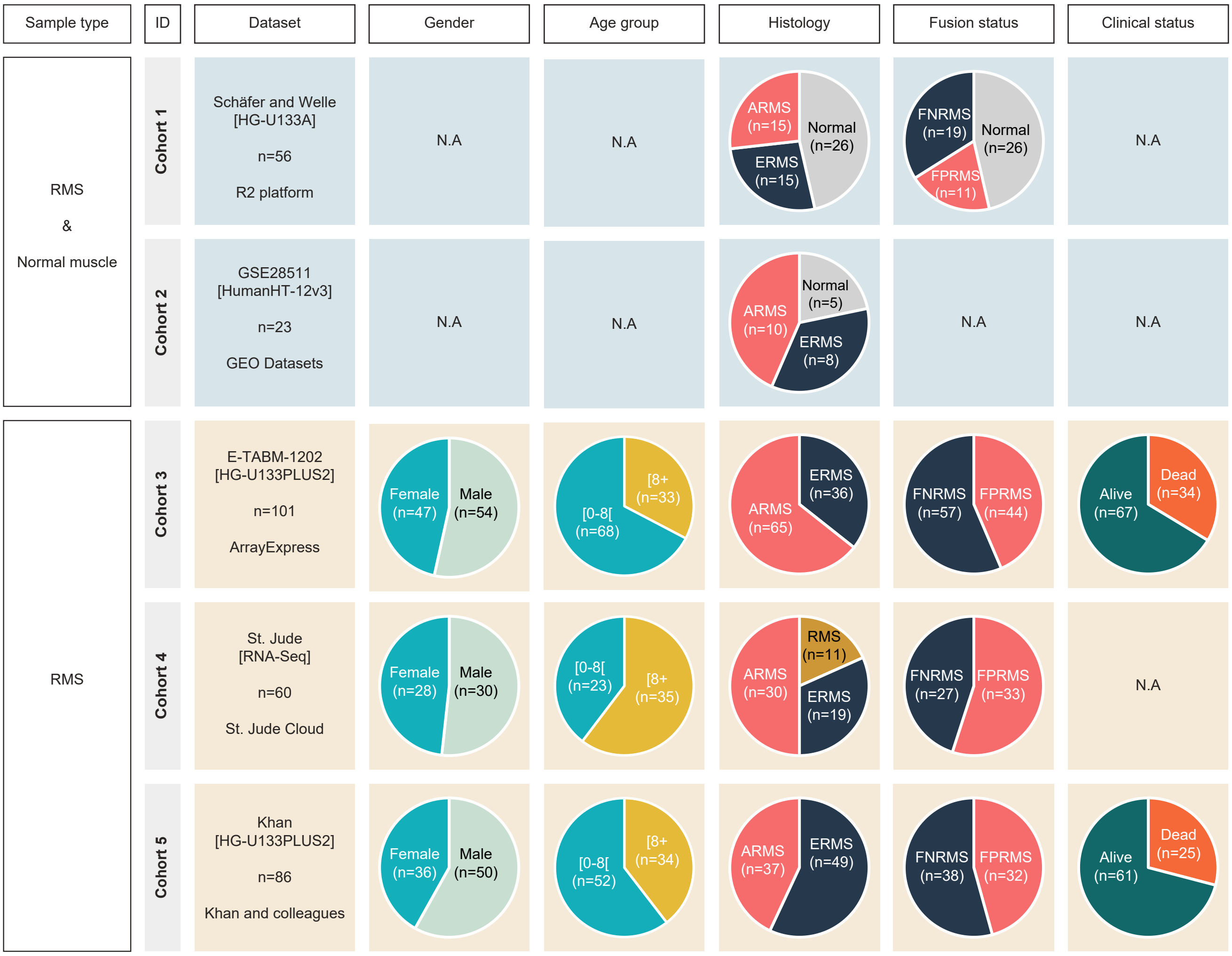

### Extended Data Figure 2

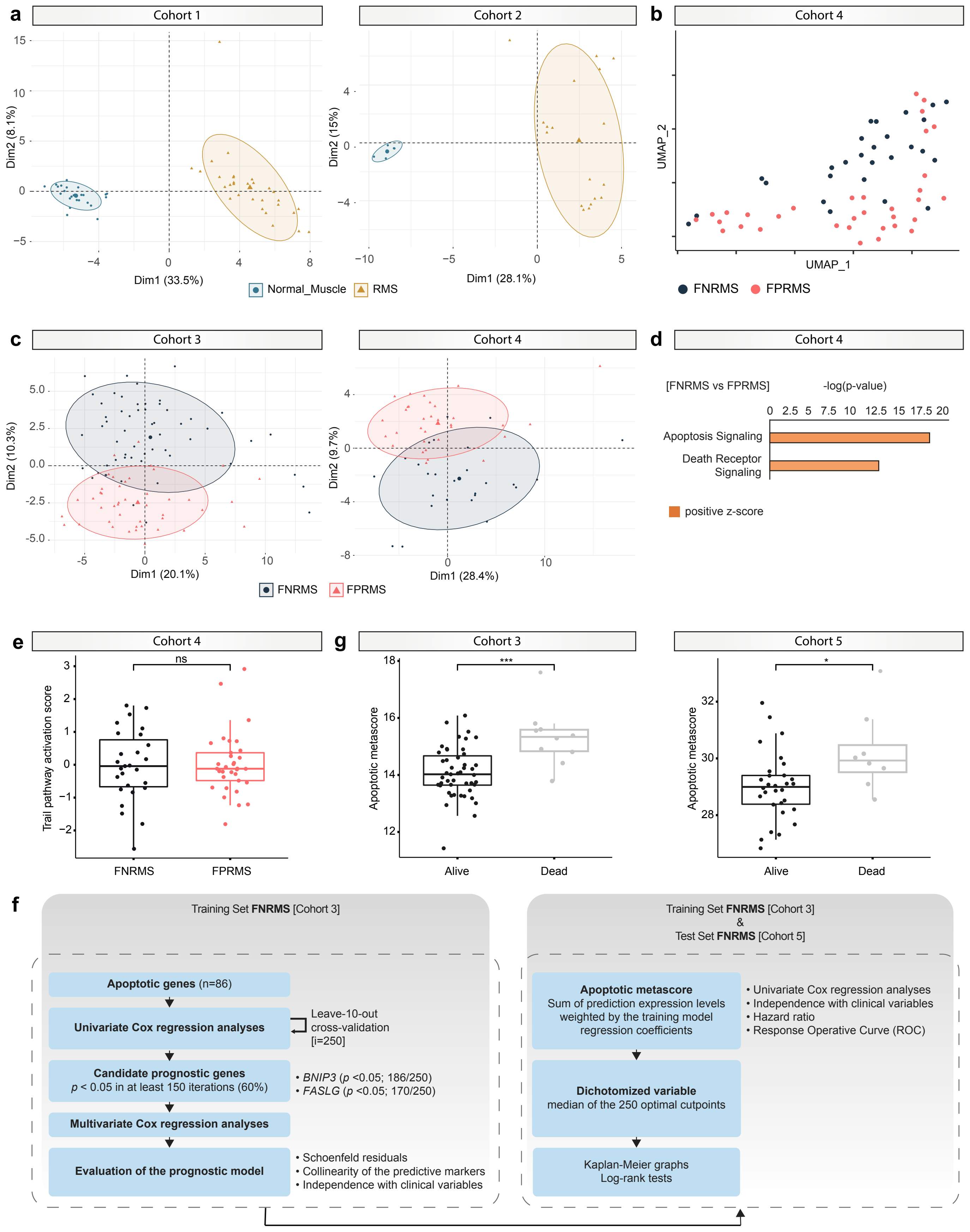

### Extended Data Figure 3

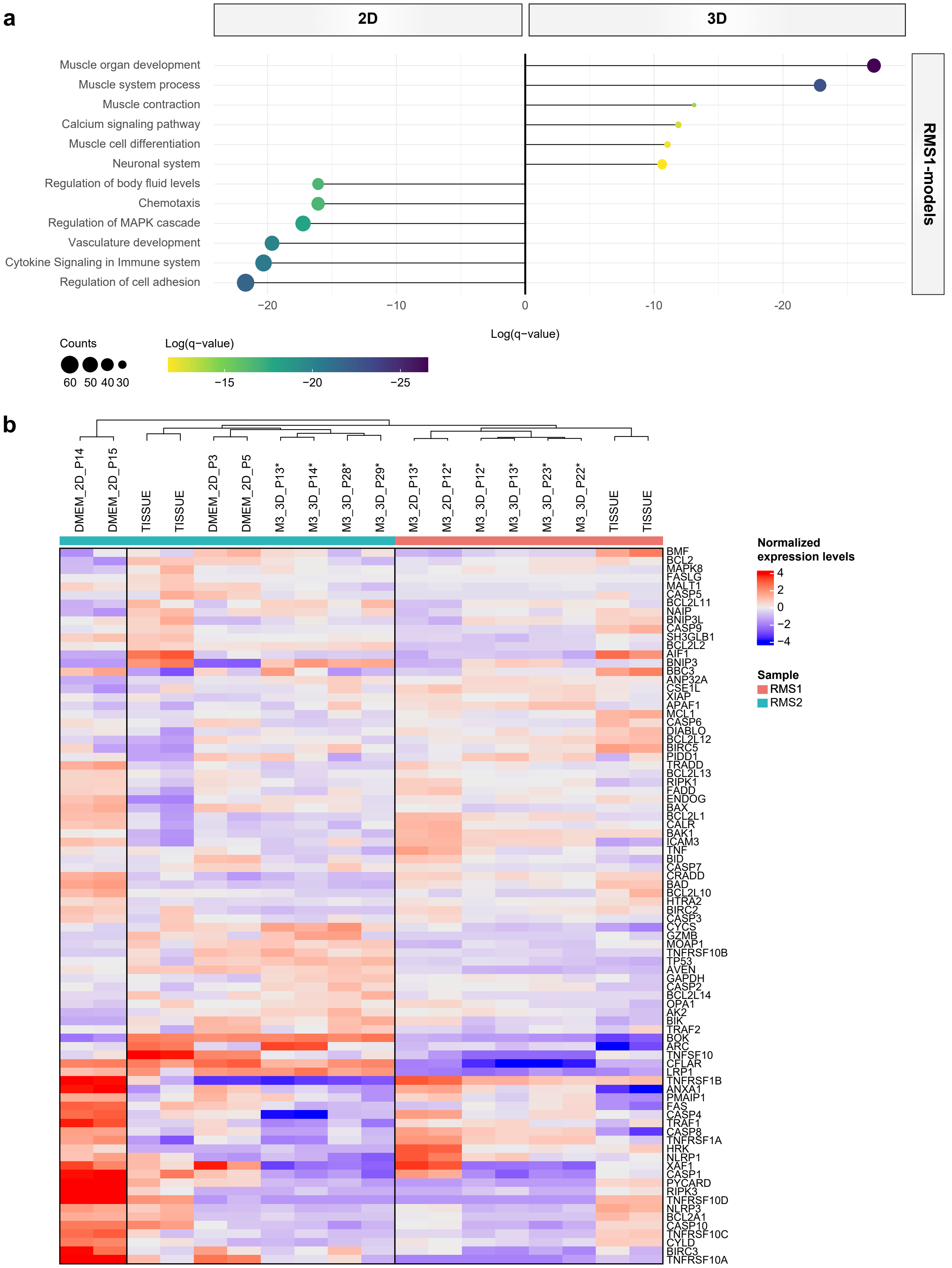

### Extended Data Figure 4

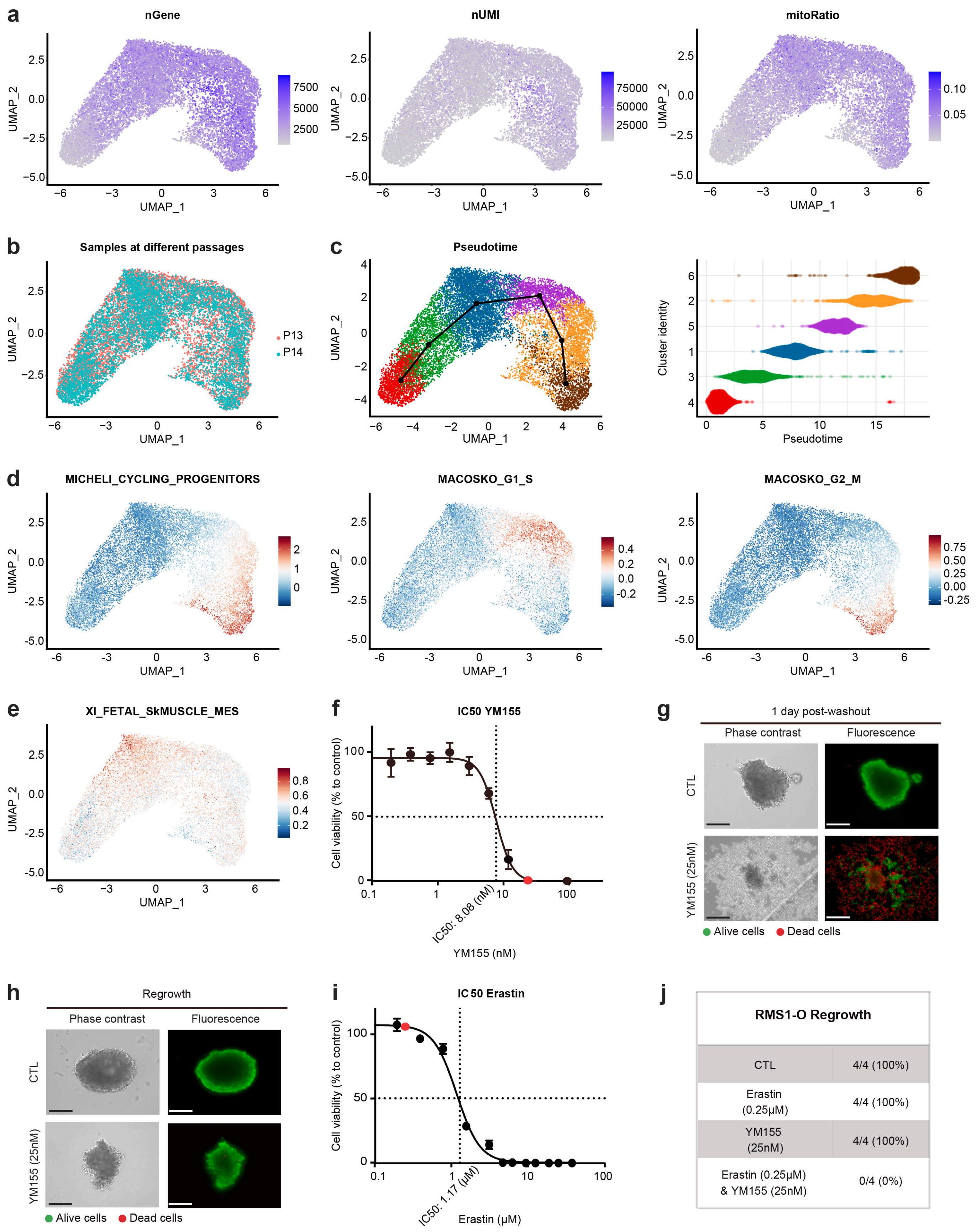


| RMS1-O Regrowth | |
|---|---|
| CTL | 4/4 (100%) |
| Erastin (0.25μM) | 4/4 (100%) |
| YM155 (25nM) | 4/4 (100%) |
| Erastin (0.25μM) & YM155 (25nM) | 0/4 (0%) |

**Supplementary table 1.** Apoptotic genes list from the Deathbase database (http://deathbase.org/).

| hgnc_symbol | aliases | description |
|---|---|---|
| **AIF1** | AIF-1, IBA1, IRT-1, IRT1 | allograft inflammatory factor 1 |
| **AK2** | ADK2 | adenylate kinase 2 |
| **ANP32A** | C15orf1, HPPCn, I1PP2A, LANP, MAPM, PHAP1, PHAPI, PP32 | acidic nuclear phosphoprotein 32 family member A |
| **ANXA1** | ANX1, LPC1 | annexin A1 |
| **APAF1** | APAF-1, CED4 | apoptotic peptidase activating factor 1 |
| **ARC** | Arg3.1, hArc | activity regulated cytoskeleton associated protein |
| **AVEN** | PDCD12 | apoptosis and caspase activation inhibitor |
| **BAD** | BBC2, BCL2L8 | BCL2 associated agonist of cell death |
| **BAK1** | BAK, BAK-LIKE, BCL2L7, CDN1 | BCL2 antagonist/killer 1 |
| **BAX** | BCL2L4 | BCL2 associated X, apoptosis regulator |
| **BBC3** | JFY-1, JFY1, PUMA | BCL2 binding component 3 |
| **BCL2** | Bcl-2, PPP1R50 | BCL2 apoptosis regulator |
| **BCL2A1** | ACC-1, ACC-2, ACC1, ACC2, BCL2L5, BFL1, GRS, HBPA1 | BCL2 related protein A1 |
| **BCL2L1** | BCL-XL/S, BCL2L, BCLX, Bcl-X, PPP1R52 | BCL2 like 1 |
| **BCL2L10** | BCL-B, Boo, Diva, bcl2-L-10 | BCL2 like 10 |
| **BCL2L11** | BAM, BIM, BOD | BCL2 like 11 |
| **BCL2L12** | - | BCL2 like 12 |
| **BCL2L13** | BCL-RAMBO, Bcl2-L-13, MIL1 | BCL2 like 13 |
| **BCL2L14** | BCLG | BCL2 like 14 |
| **BCL2L2** | BCL-W, BCL2-L-2, BCLW, PPP1R51 | BCL2 like 2 |
| **BID** | FP497 | BH3 interacting domain death agonist |
| **BIK** | BIP1, BP4, NBK | BCL2 interacting killer |
| **BIRC2** | API1, HIAP2, Hiap-2, MIHB, RNF48, c-IAP1, cIAP1 | baculoviral IAP repeat containing 2 |
| **BIRC3** | AIP1, API2, CIAP2, HAIP1, HIAP1, IAP-1, MALT2, MIHC, RNF49, c-IAP2 | baculoviral IAP repeat containing 3 |
| **BIRC5** | API4, EPR-1 | baculoviral IAP repeat containing 5 |
| **BMF** | - | Bcl2 modifying factor |
| **BNIP3** | HABON, NIP3 | BCL2 interacting protein 3 |
| **BNIP3L** | BNIP3a, NIX | BCL2 interacting protein 3 like |
| **BOK** | BCL2L9, BOKL | BCL2 family apoptosis regulator BOK |
| **CALR** | CRT, HEL-S-99n, RO, SSA, cC1qR | calreticulin |
| **CASP1** | ICE, IL1BC, P45 | caspase 1 |
| **CASP10** | ALPS2, FLICE-2, FLICE2, MCH4 | caspase 10 |
| **CASP12** | CASP-12, CASP12P1 | caspase 12 (gene/pseudogene) |
| **CASP14** | ARCI12 | caspase 14 |
| **CASP2** | CASP-2, ICH1, NEDD-2, NEDD2, PPP1R57 | caspase 2 |
| **CASP3** | CPP32, CPP32B, SCA-1 | caspase 3 |
| **CASP4** | ICE(rel)II, ICEREL-II, ICH-2, Mih1, Mih1/TX, TX | caspase 4 |
| **CASP5** | ICE(rel)III, ICEREL-III, ICH-3 | caspase 5 |
| **CASP6** | CSP-6, MCH2 | caspase 6 |
| **CASP7** | CASP-7, CMH-1, ICE-LAP3, LICE2, MCH3 | caspase 7 |
| **CASP8** | ALPS2B, CAP4, Casp-8, FLICE, MACH, MCH5 | caspase 8 |
| **CASP9** | APAF-3, APAF3, ICE-LAP6, MCH6, PPP1R56 | caspase 9 |
| **CFLAR** | CASH, CASP8AP1, CLARP, Casper, FLAME, FLAME-1, FLAME1, FLIP, I-FLICE, MRIT, c-FLIP, c-FLIPL, c-FLIPR, c-FLIPS, cFLIP | CASP8 and FADD like apoptosis regulator |
| **CRADD** | MRT34, RAIDD | CASP2 and RIPK1 domain containing adaptor with death domain |
| **CSE1L** | CAS, CSE1, XPO2 | chromosome segregation 1 like |
| **CYCS** | CYC, HCS, THC4 | cytochrome c, somatic |
| **CYLD** | BRSS, CDMT, CYLD1, CYLDI, EAC, FTDALS8, MFT, MFT1, SBS, TEM, USPL2 | CYLD lysine 63 deubiquitinase |
| **DIABLO** | DFNA64, SMAC | diablo IAP-binding mitochondrial protein |
| **ENDOG** | - | endonuclease G |
| **FADD** | GIG3, IMD90, MORT1 | Fas associated via death domain |
| **FAS** | ALPS1A, APO-1, APT1, CD95, FAS1, FASTM, TNFRSF6 | Fas cell surface death receptor |
| **FASLG** | ALPS1B, APT1LG1, APTL, CD178, CD95-L, CD95L, FASL, TNFSF6, TNLG1A | Fas ligand |
| **GAPDH** | G3PD, GAPD, HEL-S-162eP | glyceraldehyde-3-phosphate dehydrogenase |
| **GZMB** | C11, CCPI, CGL-1, CGL1, CSP-B, CSPB, CTLA1, CTSGL1, HLP, SECT | granzyme B |
| **HRK** | DP5, HARAKIRI | harakiri, BCL2 interacting protein |
| **HTRA2** | MGCA8, OMI, PARK13, PRSS25 | HtrA serine peptidase 2 |
| **ICAM3** | CD50, CDW50, ICAM-R | intercellular adhesion molecule 3 |
| **LRP1** | A2MR, APOER, APR, CD91, IGFBP-3R, IGFBP3R, IGFBP3R1, KPA, LRP, LRP1A, TGFBR5 | LDL receptor related protein 1 |
| **MALT1** | IMD12, MLT, MLT1, PCASP1 | MALT1 paracaspase |
| **MAPK8** | JNK, JNK-46, JNK1, JNK1A2, JNK21B1/2, PRKM8, SAPK1, SAPK1c | mitogen-activated protein kinase 8 |
| **MCL1** | BCL2L3, EAT, MCL1-ES, MCL1L, MCL1S, Mcl-1, TM, bcl2-L-3, mcl1/EAT | MCL1 apoptosis regulator, BCL2 family member |
| **MOAP1** | MAP-1, PNMA4 | modulator of apoptosis 1 |
| **NAIP** | BIRC1, NLRB1, psiNAIP | NLR family apoptosis inhibitory protein |
| **NLRP1** | AIADK, CARD7, CIDED, CLR17.1, DEFCAP, DEFCAP-L/S, JRRP, MSPC, NAC, NALP1, PP1044, SLEV1, VAMAS1 | NLR family pyrin domain containing 1 |
| **NLRP3** | AGTAVPRL, AII, AVP, C1orf7, CIAS1, CLR1.1, DFNA34, FCAS, FCAS1, FCU, KEFH, MWS, NALP3, PYPAF1 | NLR family pyrin domain containing 3 |
| **OPA1** | BERHS, MGM1, MTDPS14, NPG, NTG, largeG | OPA1 mitochondrial dynamin like GTPase |
| **PIDD1** | LRDD, MRT75, PIDD | p53-induced death domain protein 1 |
| **PMAIP1** | APR, NOXA | phorbol-12-myristate-13-acetate-induced protein 1 |
| **PYCARD** | ASC, CARD5, TMS, TMS-1, TMS1 | PYD and CARD domain containing |
| **RIPK1** | AIEFL, IMD57, RIP, RIP-1, RIP1 | receptor interacting serine/threonine kinase 1 |
| **RIPK3** | RIP3 | receptor interacting serine/threonine kinase 3 |
| **SH3GLB1** | Bif-1, CGI-61, PPP1R70, dJ612B15.2 | SH3 domain containing GRB2 like, endophilin B1 |
| **TNF** | DIF, TNF-alpha, TNFA, TNFSF2, TNLG1F | tumor necrosis factor |
| **TNFRSF10A** | APO2, CD261, DR4, TRAILR-1, TRAILR1 | TNF receptor superfamily member 10a |
| **TNFRSF10B** | CD262, DR5, KILLER, KILLER/DR5, TRAIL-R2, TRAILR2, TRICK2, TRICK2A, TRICK2B, TRICKB, ZTNFR9 | TNF receptor superfamily member 10b |
| **TNFRSF10C** | CD263, DCR1, DCR1-TNFR, LIT, TRAIL-R3, TRAILR3, TRID | TNF receptor superfamily member 10c |
| **TNFRSF10D** | CD264, DCR2, TRAIL-R4, TRAILR4, TRUNDD | TNF receptor superfamily member 10d |
| **TNFRSF1A** | CD120a, FPF, TBP1, TNF-R, TNF-R-I, TNF-R55, TNFAR, TNFR1, TNFR55, TNFR60, p55, p55-R, p60 | TNF receptor superfamily member 1A |
| **TNFRSF1B** | CD120b, TBPII, TNF-R-II, TNF-R75, TNFBR, TNFR1B, TNFR2, TNFR80, p75, p75TNFR | TNF receptor superfamily member 1B |
| **TNFSF10** | APO2L, Apo-2L, CD253, TL2, TNLG6A, TRAIL | TNF superfamily member 10 |
| **TP53** | BCC7, BMFS5, LFS1, P53, TRP53 | tumor protein p53 |
| **TRADD** | Hs.89862 | TNFRSF1A associated via death domain |
| **TRAF1** | EBI6, MGC:10353 | TNF receptor associated factor 1 |
| **TRAF2** | MGC:45012, RNF117, TRAP, TRAP3 | TNF receptor associated factor 2 |
| **XAF1** | BIRC4BP, HSXIAPAF1, XIAPAF1 | XIAP associated factor 1 |
| **XIAP** | API3, BIRC4, IAP-3, ILP1, MIHA, XLP2, hIAP-3, hIAP3 | X-linked inhibitor of apoptosis |

**Supplementary table 2.** Differential gene expression analysis of apoptotic genes in cohorts 1 to 5.

| | | | Cohort 1 | [RMS vs Normal Muscle] | | | [FNRMS vs Normal Muscle] | | |
|---|---|---|---|---|---|---|---|---|---|
| hgnc_symbol | entrez_id | apoptotic_status | probeset_id | log2FC | p_value | fdr | logFC | P.Value | adj.P.Val |
| **AIF1** | 199 | Pro-apoptotic | 215051_x_at | 2,031026506 | 4,84654E-08 | 1,2601E-07 | 1,982761475 | 1,45589E-06 | 3,24455E-06 |
| **AK2** | 204 | Pro-apoptotic | 208967_s_at | -1,247327036 | 2,08029E-12 | 1,01414E-11 | -1,29385426 | 8,06695E-10 | 3,3117E-09 |
| **ANP32A** | 8125 | Pro-apoptotic | 201051_at | 0,964645347 | 1,12445E-10 | 4,38534E-10 | 0,93853039 | 6,67271E-08 | 1,79473E-07 |
| **ANXA1** | 301 | Pro-apoptotic | 201012_at | 1,43881831 | 1,73437E-09 | 5,41123E-09 | 1,503205038 | 3,24171E-08 | 9,36495E-08 |
| **APAF1** | 317 | Pro-apoptotic | 204859_s_at | 0,734186788 | 3,28775E-08 | 8,84291E-08 | 0,864873009 | 5,67828E-09 | 2,01321E-08 |
| **ARC** | 23237 | Anti-apoptotic | 210090_at | -0,26358894 | 0,525640797 | 0,546666429 | 0,281060303 | 0,552237808 | 0,590062316 |
| **AVEN** | 57099 | Anti-apoptotic | 219366_at | -0,408687272 | 0,025123387 | 0,034993289 | -0,28952416 | 0,166621766 | 0,203070278 |
| **BAD** | 572 | Pro-apoptotic | 209364_at | -0,829647526 | 2,32355E-05 | 4,8983E-05 | -0,76794306 | 0,000560631 | 0,001093231 |
| **BAK1** | 578 | Pro-apoptotic | 203728_at | -0,208751267 | 0,266781803 | 0,315287585 | -0,16283882 | 0,443404809 | 0,487120776 |
| **BAX** | 581 | Pro-apoptotic | 211833_s_at | 2,844908981 | 3,30443E-21 | 1,28873E-19 | 2,908438782 | 5,66392E-17 | 1,10446E-15 |
| **BBC3** | 27113 | Pro-apoptotic | 211692_s_at | -0,573246834 | 0,01716381 | 0,024341403 | -0,64324376 | 0,024629925 | 0,036247815 |
| **BCL2** | 596 | Anti-apoptotic | 203685_at | -1,237441198 | 5,48546E-08 | 1,38021E-07 | -1,3197918 | 7,85748E-07 | 1,8026E-06 |
| **BCL2A1** | 597 | Anti-apoptotic | 205681_at | 0,452227302 | 0,253379164 | 0,304054997 | 0,729921094 | 0,108713802 | 0,141327943 |
| **BCL2L1** | 598 | Anti-apoptotic | 212312_at | 0,377897684 | 0,029159925 | 0,039903056 | 0,596263337 | 0,001616687 | 0,002865946 |
| **BCL2L10** | 10017 | Anti-apoptotic | 221320_at | -1,125980072 | 0,001074144 | 0,00186185 | -1,07108848 | 0,008772273 | 0,013416418 |
| **BCL2L11** | 10018 | Pro-apoptotic | 222343_at | -0,038495787 | 0,834629672 | 0,834629672 | -0,07197671 | 0,736620457 | 0,758178689 |
| **BCL2L12** | 83596 | Anti-apoptotic | | | | | | | |
| **BCL2L13** | 23786 | Pro-apoptotic | 217955_at | -1,341123962 | 6,1393E-19 | 1,19716E-17 | -1,52576429 | 1,4253E-18 | 1,11174E-16 |
| **BCL2L14** | 79370 | Pro-apoptotic | 221241_s_at | -0,591410033 | 0,002109832 | 0,003501424 | -0,63022894 | 0,002700398 | 0,004542091 |
| **BCL2L2** | 599 | Anti-apoptotic | 209311_at | 0,123898428 | 0,232840579 | 0,283774456 | 0,087672442 | 0,479657981 | 0,519629479 |
| **BID** | 637 | Pro-apoptotic | 211725_s_at | 2,530629916 | 2,63406E-14 | 1,58043E-13 | 2,577755086 | 9,90911E-11 | 4,54653E-10 |
| **BIK** | 638 | Pro-apoptotic | 205780_at | -2,325361471 | 6,01144E-19 | 1,19716E-17 | -2,40638905 | 3,5429E-15 | 3,9478E-14 |
| **BIRC2** | 329 | Anti-apoptotic | 202076_at | 0,467348641 | 2,41609E-07 | 5,88922E-07 | 0,43733259 | 1,05601E-05 | 2,28803E-05 |
| **BIRC3** | 330 | Anti-apoptotic | 210538_s_at | -0,816548261 | 0,016834691 | 0,024316776 | -0,86157878 | 0,024479279 | 0,036247815 |
| **BIRC5** | 332 | Anti-apoptotic | 202095_s_at | 2,911090338 | 1,01149E-16 | 9,86199E-16 | 2,890746263 | 4,60257E-15 | 4,4875E-14 |
| **BMF** | 90427 | Pro-apoptotic | | | | | | | |
| **BNIP3** | 664 | Pro-apoptotic | 201849_at | -0,868541141 | 5,02814E-05 | 0,000103209 | -1,22822345 | 6,68259E-07 | 1,62888E-06 |
| **BNIP3L** | 665 | Pro-apoptotic | 221478_at | 1,807014283 | 5,35497E-17 | 5,96696E-16 | 1,985530498 | 1,0815E-15 | 1,68714E-14 |
| **BOK** | 666 | Pro-apoptotic | | | | | | | |
| **CALR** | 811 | Anti-apoptotic | 212953_x_at | 0,108181098 | 0,396654232 | 0,429708752 | -0,04808842 | 0,73873821 | 0,758178689 |
| **CASP1** | 834 | Pro-apoptotic | 211366_x_at | -0,478046456 | 0,000364427 | 0,000693301 | -0,48530296 | 0,002736901 | 0,004542091 |
| **CASP10** | 843 | Pro-apoptotic | 211888_x_at | -2,133680819 | 7,69359E-13 | 4,00067E-12 | -2,36226123 | 1,2772E-11 | 7,11581E-11 |
| **CASP12** | 100506742 | Pro-apoptotic | | | | | | | |
| **CASP14** | 23581 | Pro-apoptotic | | | | | | | |
| **CASP2** | 835 | Pro-apoptotic | 208050_s_at | -0,385213319 | 0,009051621 | 0,013843656 | -0,32754017 | 0,041217623 | 0,059536567 |
| **CASP3** | 836 | Pro-apoptotic | 202763_at | 0,288007311 | 0,079658702 | 0,103556312 | 0,343678345 | 0,085511566 | 0,117015827 |
| **CASP4** | 837 | Pro-apoptotic | 209310_s_at | 0,244054219 | 0,123826833 | 0,155782145 | 0,270401654 | 0,157610346 | 0,195136619 |
| **CASP5** | 838 | Pro-apoptotic | 207500_at | -2,304153351 | 7,43088E-11 | 3,05057E-10 | -2,3595417 | 1,53893E-09 | 5,71604E-09 |
| **CASP6** | 839 | Pro-apoptotic | 211464_x_at | 0,684255325 | 8,02409E-05 | 0,00015647 | 0,761858459 | 1,16098E-05 | 2,44748E-05 |
| **CASP7** | 840 | Pro-apoptotic | 207181_s_at | -0,329654964 | 0,005422764 | 0,008459512 | -0,42670374 | 0,003041846 | 0,004942999 |
| **CASP8** | 841 | Pro-apoptotic | 213373_s_at | 0,936174544 | 0,001062711 | 0,00186185 | 1,117152059 | 0,000283558 | 0,000567116 |
| **CASP9** | 842 | Pro-apoptotic | 203984_s_at | 0,331348534 | 0,069450264 | 0,093398632 | -0,02113929 | 0,897770732 | 0,897770732 |
| **CFLAR** | 8837 | Anti-apoptotic | 211316_x_at | -0,902346306 | 4,82819E-09 | 1,345E-08 | -0,99791486 | 1,01025E-08 | 3,28331E-08 |
| **CRADD** | 8738 | Pro-apoptotic | 209833_at | -0,962257804 | 4,45431E-09 | 1,2868E-08 | -1,00605973 | 1,16823E-07 | 3,03741E-07 |
| **CSE1L** | 1434 | Anti-apoptotic | 201112_s_at | 1,174535591 | 2,464E-17 | 3,20319E-16 | 1,196755567 | 3,34037E-14 | 2,89499E-13 |
| **CYCS** | 54205 | Pro-apoptotic | 208905_at | -0,938084525 | 1,73351E-09 | 5,41123E-09 | -0,82070112 | 6,94861E-07 | 1,6424E-06 |
| **CYLD** | 1540 | Pro-apoptotic | 221905_at | 0,646968768 | 0,015529395 | 0,023294093 | 0,463274268 | 0,149835914 | 0,188503247 |
| **DIABLO** | 56616 | Pro-apoptotic | 219350_s_at | 0,389379546 | 0,005099881 | 0,008118179 | 0,445521606 | 0,001925908 | 0,003338241 |
| **ENDOG** | 2021 | Pro-apoptotic | 204824_at | -2,982837405 | 1,95376E-15 | 1,38539E-14 | -3,59922453 | 3,1618E-18 | 1,2331E-16 |
| **FADD** | 8772 | Pro-apoptotic | 202535_at | 1,282462952 | 1,14245E-06 | 2,47531E-06 | 1,280730222 | 4,59866E-05 | 9,43936E-05 |
| **FAS** | 355 | Pro-apoptotic | 204780_s_at | 2,007809699 | 1,61247E-09 | 5,41123E-09 | 2,560261309 | 1,40219E-11 | 7,29137E-11 |
| **FASLG** | 356 | Pro-apoptotic | 210865_at | -1,591215205 | 4,28215E-15 | 2,7834E-14 | -1,84517617 | 2,88417E-15 | 3,74942E-14 |
| **GAPDH** | 2597 | Pro-apoptotic | 212581_x_at | 21,54737225 | 7,36619E-24 | 5,74562E-22 | 21,50709543 | 1,29689E-17 | 3,37191E-16 |
| **GZMB** | 3002 | Pro-apoptotic | 210164_at | -0,778054064 | 0,001709076 | 0,002897998 | -0,79645237 | 0,006570119 | 0,010249385 |
| **HRK** | 8739 | Pro-apoptotic | 206864_s_at | 0,315833051 | 0,313257793 | 0,349388874 | 0,445778751 | 0,193728344 | 0,232474013 |
| **HTRA2** | 27429 | Pro-apoptotic | 203089_s_at | 1,133393431 | 1,28733E-10 | 4,78152E-10 | 1,171272499 | 2,44334E-08 | 7,62322E-08 |
| **ICAM3** | 3385 | Pro-apoptotic | 204949_at | -0,846973821 | 0,000616836 | 0,001118912 | -0,7840877 | 0,005368109 | 0,008545154 |
| **LRP1** | 4035 | Pro-apoptotic | 200785_s_at | 0,128704733 | 0,482082625 | 0,508141145 | 0,34423703 | 0,097980988 | 0,131767535 |
| **MALT1** | 10892 | Anti-apoptotic | 210018_x_at | 0,221459395 | 0,300751746 | 0,344979944 | 0,378935108 | 0,147852445 | 0,188503247 |
| **MAPK8** | 5599 | Anti-apoptotic | 210671_x_at | -1,666693466 | 1,36473E-11 | 6,26169E-11 | -1,56401139 | 1,45677E-10 | 6,31269E-10 |
| **MCL1** | 4170 | Anti-apoptotic | 200797_s_at | 0,46434139 | 0,004829684 | 0,007848236 | 0,598771322 | 0,00151981 | 0,002756864 |
| **MOAP1** | 64112 | Pro-apoptotic | 212508_at | 0,097104135 | 0,46884227 | 0,500954754 | 0,040232143 | 0,790195484 | 0,800457763 |
| **NAIP** | 4671 | Anti-apoptotic | | | | | | | |
| **NLRP1** | 22861 | Ambiguous | 211824_x_at | -1,500020436 | 6,89813E-07 | 1,58251E-06 | -1,68909331 | 2,62187E-07 | 6,59695E-07 |
| **NLRP3** | 114548 | Ambiguous | 207075_at | 1,302904795 | 1,13745E-06 | 2,47531E-06 | 1,531302953 | 5,24707E-08 | 1,46168E-07 |
| **OPA1** | 4976 | Anti-apoptotic | 212213_x_at | -0,259784623 | 0,098010528 | 0,125324938 | -0,34036906 | 0,07531175 | 0,104898508 |
| **PIDD1** | 55367 | Pro-apoptotic | 221640_s_at | -0,108803911 | 0,689243736 | 0,698194954 | -0,11248332 | 0,722030066 | 0,758178689 |
| **PMAIP1** | 5366 | Pro-apoptotic | 204285_s_at | 3,937781114 | 2,88342E-14 | 1,60647E-13 | 4,338728812 | 1,95952E-12 | 1,17571E-11 |
| **PYCARD** | 29108 | Pro-apoptotic | 221666_s_at | 2,385939329 | 2,05601E-16 | 1,60369E-15 | 2,462352112 | 4,93129E-13 | 3,49673E-12 |
| **RIPK1** | 8737 | Pro-apoptotic | 209941_at | -0,743910187 | 0,000510792 | 0,000948615 | -0,67898535 | 0,00116458 | 0,002162791 |
| **RIPK3** | 11035 | Pro-apoptotic | | | | | | | |
| **SH3GLB1** | 51100 | Pro-apoptotic | 209090_s_at | -0,691992132 | 6,51344E-07 | 1,53954E-06 | -0,9771047 | 5,31459E-11 | 2,59086E-10 |
| **TNF** | 7124 | Pro-apoptotic | 207113_s_at | -0,149902862 | 0,568582139 | 0,583544827 | -0,23667177 | 0,432334414 | 0,481744061 |
| **TNFRSF10A** | 8797 | Pro-apoptotic | | | | | | | |
| **TNFRSF10B** | 8795 | Pro-apoptotic | 209295_at | 2,951197557 | 1,6057E-16 | 1,39161E-15 | 3,036129733 | 6,02706E-13 | 3,91759E-12 |
| **TNFRSF10C** | 8794 | Pro-apoptotic | 206222_at | -0,386925002 | 0,071320613 | 0,094288269 | -0,42043736 | 0,102027143 | 0,134883341 |
| **TNFRSF10D** | 8793 | Pro-apoptotic | 210654_at | -1,349989347 | 6,52948E-05 | 0,00013059 | -1,2796849 | 0,001146069 | 0,002162791 |
| **TNFRSF1A** | 7132 | Pro-apoptotic | 207643_s_at | -0,436442361 | 0,016653487 | 0,024316776 | -0,23183545 | 0,250194158 | 0,291270811 |
| **TNFRSF1B** | 7133 | Pro-apoptotic | 203508_at | -0,221435362 | 0,157693316 | 0,195239344 | -0,34474707 | 0,071937581 | 0,102020569 |
| **TNFSF10** | 8743 | Pro-apoptotic | 202688_at | -1,801014253 | 2,56865E-10 | 9,10702E-10 | -1,91747187 | 8,01167E-09 | 2,717E-08 |
| **TP53** | 7157 | Pro-apoptotic | 201746_at | 0,161760933 | 0,325408981 | 0,357491556 | 0,220918063 | 0,256650101 | 0,294392763 |
| **TRADD** | 8717 | Pro-apoptotic | 205641_s_at | 2,397774708 | 4,91679E-18 | 7,6702E-17 | 2,152458061 | 1,11491E-13 | 8,69627E-13 |
| **TRAF1** | 7185 | Pro-apoptotic | 205599_at | -1,137441168 | 3,6494E-09 | 1,09482E-08 | -1,30156663 | 2,70142E-08 | 8,10425E-08 |
| **TRAF2** | 7186 | Pro-apoptotic | 204413_at | 1,634174112 | 4,32245E-11 | 1,87306E-10 | 1,688318571 | 1,2528E-09 | 4,88592E-09 |
| **XAF1** | 54739 | Pro-apoptotic | 206133_at | -0,334276452 | 0,313554118 | 0,349388874 | -0,30609643 | 0,419412577 | 0,474118566 |
| **XIAP** | 331 | Anti-apoptotic | 206536_s_at | -0,15222604 | 0,30000755 | 0,344979944 | -0,20138332 | 0,23836965 | 0,281709586 |

| | Cohort 2 | [RMS vs Normal Muscle] | | | [ERMS vs Normal Muscle] | | |
|---|---|---|---|---|---|---|---|
| hgnc_symbol | probeset_id | log2FC | p_value | fdr | logFC | P.Value | adj.P.Val |
| **AIF1** | ILMN_1792473 | 0,205760675 | 0,34952312 | 0,469671688 | 0,351208004 | 0,057341387 | 0,117413317 |
| **AK2** | ILMN_1716053 | -1,111281009 | 1,3926E-11 | 3,99214E-10 | -1,1691095 | 4,87249E-10 | 4,19034E-08 |
| **ANP32A** | ILMN_1807945 | 0,103840827 | 0,15955848 | 0,285875616 | 0,047889328 | 0,458051134 | 0,596854508 |
| **ANXA1** | ILMN_2184184 | -1,042450946 | 0,06049903 | 0,130072915 | -1,28675132 | 0,022462826 | 0,060368845 |
| **APAF1** | ILMN_1659463 | 1,163570133 | 8,0721E-11 | 1,73551E-09 | 1,194704919 | 4,55509E-08 | 4,35264E-07 |
| **ARC** | ILMN_1711120 | 0,141000986 | 0,19002194 | 0,333507891 | 0,192098633 | 0,081557667 | 0,152477377 |
| **AVEN** | ILMN_1710216 | -0,455266942 | 2,6998E-05 | 0,000122202 | -0,52729103 | 4,99187E-05 | 0,000195137 |
| **BAD** | ILMN_1738652 | -0,275574978 | 0,14596606 | 0,272893066 | 0,072257094 | 0,612284829 | 0,71157426 |
| **BAK1** | ILMN_1805990 | 0,145784158 | 0,23836594 | 0,394740372 | 0,104586405 | 0,538581404 | 0,661614096 |
| **BAX** | ILMN_2321064 | 1,655511602 | 9,5671E-05 | 0,000373986 | 2,13212428 | 2,00033E-06 | 1,07518E-05 |
| **BBC3** | ILMN_1729645 | -0,153272799 | 0,05010205 | 0,110481452 | -0,14641252 | 0,104326387 | 0,183103455 |
| **BCL2** | ILMN_2246956 | -0,919098473 | 6,5694E-08 | 5,64968E-07 | -0,85980694 | 2,81686E-07 | 2,20228E-06 |
| **BCL2A1** | ILMN_1769229 | 0,146754523 | 0,01010625 | 0,027160551 | 0,117294978 | 0,072378546 | 0,141467158 |
| **BCL2L1** | ILMN_1654118 | 0,146262615 | 0,5148591 | 0,651145337 | 0,460628688 | 0,031132294 | 0,077919582 |
| **BCL2L10** | ILMN_1749096 | 0,146456088 | 0,08632948 | 0,176769884 | 0,125789866 | 0,140545541 | 0,24173833 |
| **BCL2L11** | ILMN_2359627 | 0,171574217 | 0,0729771 | 0,153073913 | 0,134154645 | 0,101610846 | 0,182052765 |
| **BCL2L12** | ILMN_2396982 | 0,990775419 | 0,00064124 | 0,00212101 | 0,691463323 | 0,036646817 | 0,087545174 |
| **BCL2L13** | ILMN_2181445 | -1,396282756 | 1,0179E-05 | 5,4714E-05 | -1,85425697 | 6,89646E-07 | 4,2364E-06 |
| **BCL2L14** | ILMN_1664235 | -0,071017847 | 0,33970863 | 0,469671688 | -0,06882113 | 0,422675325 | 0,581746986 |
| **BCL2L2** | ILMN_1746561 | -0,100915512 | 0,34480594 | 0,469671688 | -0,21383215 | 0,031711458 | 0,077919582 |
| **BID** | ILMN_1763386 | 0,554717397 | 0,00058702 | 0,002019338 | 0,381156133 | 0,03846814 | 0,089412432 |
| **BIK** | ILMN_1770505 | 0,027334623 | 0,77547623 | 0,844189311 | 0,025419125 | 0,823050956 | 0,877519507 |
| **BIRC2** | ILMN_2182704 | 0,079178713 | 0,60999868 | 0,738871639 | 0,110197576 | 0,567952845 | 0,669095133 |
| **BIRC3** | ILMN_2405684 | 0,984499904 | 0,03263333 | 0,073854387 | 1,045156395 | 0,046233327 | 0,096977222 |
| **BIRC5** | ILMN_2349459 | 3,522231275 | 3,2785E-13 | 2,81947E-11 | 3,500773347 | 2,96964E-08 | 3,19236E-07 |
| **BMF** | ILMN_2308338 | 1,053663966 | 0,00234533 | 0,006955111 | 0,903346502 | 0,021584216 | 0,059878793 |
| **BNIP3** | ILMN_1724658 | -1,657674176 | 1,3884E-06 | 7,96004E-06 | -1,79418209 | 4,83027E-05 | 0,000195137 |
| **BNIP3L** | ILMN_1718961 | 0,858727267 | 0,02667835 | 0,062009148 | 1,009903974 | 0,042511426 | 0,093267379 |
| **BOK** | ILMN_1730032 | 1,298553489 | 3,201E-07 | 2,11757E-06 | 1,217664813 | 5,09059E-07 | 3,36762E-06 |
| **CALR** | ILMN_1736256 | 1,39728055 | 1,2435E-05 | 6,29046E-05 | 1,571326259 | 5,13677E-06 | 2,32507E-05 |
| **CASP1** | ILMN_2326509 | -0,027301692 | 0,85854863 | 0,90042905 | -0,03975453 | 0,686946726 | 0,777334453 |
| **CASP10** | ILMN_1797578 | 0,079000089 | 0,32585973 | 0,467065614 | 0,114816995 | 0,258182564 | 0,385619285 |
| **CASP12** | ILMN_2132119 | -0,026369179 | 0,76229545 | 0,840479593 | 0,02241072 | 0,826500931 | 0,877519507 |
| **CASP14** | ILMN_1739513 | 0,085275523 | 0,26025107 | 0,414473923 | 0,121064303 | 0,181508044 | 0,289068366 |
| **CASP2** | ILMN_2410540 | 1,564266479 | 1,8532E-10 | 3,18747E-09 | 1,750077584 | 4,1778E-09 | 8,89752E-08 |
| **CASP3** | ILMN_2388155 | 1,075413489 | 0,01624171 | 0,039908199 | 1,637187286 | 0,002287273 | 0,007025196 |
| **CASP4** | ILMN_1678454 | -0,106926246 | 0,72108827 | 0,838021503 | -0,42773101 | 0,076392074 | 0,145993742 |
| **CASP5** | ILMN_1722158 | 0,012265944 | 0,87776321 | 0,909489595 | 0,065235061 | 0,518864566 | 0,646700764 |
| **CASP6** | ILMN_1659350 | 0,051568821 | 0,3162121 | 0,467065614 | 0,075059561 | 0,271114675 | 0,395184103 |
| **CASP7** | ILMN_2373763 | -0,24912628 | 0,08881307 | 0,177626143 | -0,21356116 | 0,257131087 | 0,385619285 |
| **CASP8** | ILMN_1669565 | 0,048174583 | 0,63531793 | 0,758851975 | 0,145394833 | 0,149445494 | 0,252006128 |
| **CASP9** | ILMN_1718070 | 0,081207433 | 0,74209916 | 0,839743787 | -0,29591997 | 0,068592979 | 0,137185958 |
| **CFLAR** | ILMN_1789830 | -1,533344027 | 9,3856E-05 | 0,000373986 | -1,93787053 | 0,000152708 | 0,000547204 |
| **CRADD** | ILMN_1690282 | -1,52979349 | 2,4421E-12 | 1,05011E-10 | -1,57574261 | 6,20757E-09 | 8,89752E-08 |
| **CSE1L** | ILMN_1706238 | 1,166250487 | 1,5432E-07 | 1,20651E-06 | 1,075236141 | 9,3425E-08 | 8,03455E-07 |
| **CYCS** | ILMN_1730416 | -1,207441365 | 3,1755E-05 | 0,000136548 | -1,45225885 | 9,92684E-07 | 5,69139E-06 |
| **CYLD** | ILMN_1775508 | -0,506853417 | 0,00021355 | 0,000798506 | -0,51018632 | 0,000957681 | 0,003167715 |
| **DIABLO** | ILMN_2310589 | 0,633284834 | 0,0002953 | 0,001058175 | 0,757824439 | 0,000272606 | 0,000937764 |
| **ENDOG** | ILMN_1722309 | -1,843913079 | 2,0482E-08 | 1,95718E-07 | -1,78104972 | 1,83799E-05 | 7,90334E-05 |
| **FADD** | ILMN_1758658 | -0,028028228 | 0,85640918 | 0,90042905 | 0,035521013 | 0,8016722 | 0,872706446 |
| **FAS** | ILMN_2319077 | -0,093223264 | 0,27783998 | 0,426682828 | -0,09154362 | 0,402562672 | 0,567547373 |
| **FASLG** | ILMN_1781824 | -0,038280498 | 0,58914758 | 0,725695395 | -0,06642685 | 0,470066257 | 0,598615578 |
| **GAPDH** | ILMN_2038778 | -1,694765656 | 3,8918E-09 | 4,78139E-08 | -1,73312933 | 6,04317E-09 | 8,89752E-08 |
| **GZMB** | ILMN_2109489 | 0,018109201 | 0,93512682 | 0,957391746 | -0,06634738 | 0,550595842 | 0,661614096 |
| **HRK** | ILMN_2193706 | -0,061544784 | 0,73420468 | 0,839743787 | 0,243946253 | 0,196558928 | 0,307346687 |
| **HTRA2** | ILMN_1747204 | 0,248678943 | 0,21713647 | 0,373474736 | 0,241999322 | 0,28908903 | 0,414360943 |
| **ICAM3** | ILMN_2212763 | -0,792103054 | 0,01522516 | 0,038510698 | -0,91395609 | 0,001506954 | 0,004799926 |
| **LRP1** | ILMN_1669772 | 0,303378697 | 0,15461869 | 0,282919296 | 0,331448093 | 0,26006882 | 0,385619285 |
| **MALT1** | ILMN_1730986 | -0,046281581 | 0,81530716 | 0,876455197 | -0,08731272 | 0,701582424 | 0,783585565 |
| **MAPK8** | ILMN_2364574 | 0,034769387 | 0,5906823 | 0,725695395 | -0,0041194 | 0,96377739 | 0,96377739 |
| **MCL1** | ILMN_1803988 | 0,348676197 | 0,0096894 | 0,026880278 | 0,306522119 | 0,043380176 | 0,093267379 |
| **MOAP1** | ILMN_1804988 | -1,203191767 | 5,4424E-10 | 7,80076E-09 | -1,35423456 | 4,4473E-09 | 8,89752E-08 |
| **NAIP** | ILMN_2345812 | -0,055462555 | 0,3904027 | 0,508706551 | -0,06370568 | 0,446493212 | 0,590744866 |
| **NLRP1** | ILMN_1810045 | -0,001936657 | 0,98167242 | 0,981672424 | 0,025444626 | 0,798442784 | 0,872706446 |
| **NLRP3** | ILMN_1712026 | 0,586654415 | 0,09918618 | 0,193863899 | -0,00700353 | 0,949854933 | 0,961029697 |
| **OPA1** | ILMN_1729775 | -0,726913662 | 1,9531E-08 | 1,95718E-07 | -0,72803271 | 2,98042E-06 | 1,50774E-05 |
| **PIDD1** | ILMN_2400352 | 1,121005034 | 0,00069426 | 0,002211349 | 1,535969437 | 5,03021E-09 | 8,89752E-08 |
| **PMAIP1** | ILMN_2098446 | 0,407055515 | 0,00345109 | 0,00989313 | 0,345697901 | 0,039959958 | 0,090435695 |
| **PYCARD** | ILMN_1698766 | 0,123944649 | 0,48242868 | 0,619236812 | -0,01670594 | 0,917760311 | 0,946923829 |
| **RIPK1** | ILMN_2119535 | 0,416630601 | 0,00156696 | 0,004812815 | 0,62309531 | 0,000107446 | 0,000401756 |
| **RIPK3** | ILMN_1763763 | 0,093746909 | 0,31413308 | 0,467065614 | 0,085763761 | 0,473323945 | 0,598615578 |
| **SH3GLB1** | ILMN_1766045 | -1,320728884 | 6,9992E-07 | 4,29953E-06 | -1,47478929 | 5,08081E-07 | 3,36762E-06 |
| **TNF** | ILMN_1728106 | -0,135421784 | 0,12183281 | 0,232836041 | -0,15622582 | 0,158144257 | 0,26154627 |
| **TNFRSF10A** | ILMN_1721316 | -0,023455641 | 0,76200897 | 0,840479593 | -0,05959287 | 0,435735771 | 0,585519943 |
| **TNFRSF10B** | ILMN_1699265 | 2,035892174 | 2,8583E-07 | 2,04846E-06 | 2,370446477 | 3,92409E-06 | 1,87484E-05 |
| **TNFRSF10C** | ILMN_1672114 | -0,121007819 | 0,2473077 | 0,40129174 | -0,07327078 | 0,553909476 | 0,661614096 |
| **TNFRSF10D** | ILMN_1666022 | -0,002702617 | 0,96228852 | 0,973609557 | 0,005469212 | 0,924902344 | 0,946923829 |
| **TNFRSF1A** | ILMN_1685005 | -0,318384228 | 0,32396828 | 0,467065614 | -0,17750107 | 0,671350052 | 0,769814727 |
| **TNFRSF1B** | ILMN_1764788 | -0,079287581 | 0,64868488 | 0,764204102 | -0,26705545 | 0,028040021 | 0,073073995 |
| **TNFSF10** | ILMN_1801307 | -1,424103565 | 2,0749E-05 | 9,91358E-05 | -1,63008329 | 1,02425E-08 | 1,25837E-07 |
| **TP53** | ILMN_1779356 | 0,259271811 | 0,01336961 | 0,03484201 | 0,307896202 | 0,00289545 | 0,008586507 |
| **TRADD** | ILMN_1765851 | 0,139617063 | 0,37699486 | 0,498793204 | -0,01861374 | 0,892525156 | 0,936062968 |
| **TRAF1** | ILMN_1698218 | 0,102550333 | 0,33412418 | 0,469671688 | 0,122701342 | 0,180032659 | 0,289068366 |
| **TRAF2** | ILMN_1691487 | 0,551575326 | 0,01930585 | 0,046119532 | 0,749111168 | 0,007260392 | 0,020813123 |
| **XAF1** | ILMN_1742618 | -0,31395171 | 0,23868022 | 0,394740372 | -0,18522115 | 0,42616349 | 0,581746986 |
| **XIAP** | ILMN_1815361 | -0,112060477 | 0,26944973 | 0,421321402 | -0,18228358 | 0,090585772 | 0,165752688 |

| | Cohort 3 [FNRMS vs FPRMS] | | | | Cohort 4 [FNRMS vs FPRMS] | | | Cohort 5 [FNRMS vs FPRMS] | | | |
|---|---|---|---|---|---|---|---|---|---|---|---|
| hgnc_symbol | probeset_id | log2FC | p_value | fdr | log2FC | p_value | fdr | probeset_id | log2FC | p_value | fdr |
| **AIF1** | 215051_x_at | 0,084423646 | 0,594225486 | 0,740628866 | 0,978536035 | 0,002232647 | 0,009488752 | 215051_x_at | 0,001055921 | 0,991335402 | 0,99211373 |
| **AK2** | 208967_s_at | 0,263712919 | 0,002016172 | 0,008669539 | -0,018284409 | 0,879501043 | 0,934469858 | 208967_s_at | -0,029463816 | 0,731094001 | 0,881541703 |
| **ANP32A** | 201051_at | 0,077963654 | 0,387946966 | 0,556057318 | 0,457023418 | 0,007064836 | 0,021446824 | 201051_at | -0,231383224 | 0,108979943 | 0,382017004 |
| **ANXA1** | 201012_at | 0,019204658 | 0,953750908 | 0,976459263 | 0,564109618 | 0,061129236 | 0,120836861 | 201012_at | 0,209194079 | 0,211171025 | 0,540322942 |
| **APAF1** | 204859_s_at | 0,265372359 | 0,008719892 | 0,02585899 | 0,360950613 | 0,017982642 | 0,050950818 | 211554_s_at | 0,067449013 | 0,299786741 | 0,6766615 |
| **ARC** | 210090_at | 0,042417979 | 0,664751549 | 0,791156151 | -1,49884476 | 0,032798851 | 0,079654352 | 210090_at | -0,000574013 | 0,99211373 | 0,99211373 |
| **AVEN** | 219366_at | 0,148034244 | 0,182499621 | 0,320305457 | 0,390554283 | 0,005219396 | 0,018323348 | 219366_at | -0,020148026 | 0,757402171 | 0,881541703 |
| **BAD** | 209364_at | 0,239807709 | 0,004955392 | 0,016390913 | 0,349409325 | 0,04616804 | 0,103270616 | 209364_at | 0,063939145 | 0,240084331 | 0,574747339 |
| **BAK1** | 203728_at | 0,187745981 | 0,00166694 | 0,007964271 | -0,192847862 | 0,070417184 | 0,136033197 | 203728_at | -0,020902961 | 0,785639519 | 0,899500319 |
| **BAX** | 208478_s_at | 0,138602352 | 0,238124073 | 0,386390006 | 0,163607517 | 0,209142438 | 0,362798107 | 211833_s_at | -0,035106908 | 0,603599704 | 0,803854064 |
| **BBC3** | 211692_s_at | 0,083340581 | 0,17390995 | 0,311588661 | 0,194718052 | 0,319628571 | 0,485150509 | 211692_s_at | 0,097297697 | 0,088632292 | 0,377527855 |
| **BCL2** | 203685_at | -0,121978491 | 0,550248075 | 0,716989916 | -0,153461994 | 0,395799577 | 0,570219729 | 203684_s_at | 0,050638158 | 0,447090809 | 0,768184772 |
| **BCL2A1** | 205681_at | 0,214467224 | 0,539214619 | 0,713422419 | 1,512971406 | 0,86916582 | 0,934469858 | 205681_at | 0,017060855 | 0,835871405 | 0,930054098 |
| **BCL2L1** | 212312_at | 0,328170507 | 0,00049929 | 0,002683682 | -0,156089395 | 0,000361315 | 0,002559314 | 212312_at | 0,003167763 | 0,950943894 | 0,984776852 |
| **BCL2L10** | 221320_at | 0,020788475 | 0,575582317 | 0,727942342 | -1,180879636 | 0,001244953 | 0,006224767 | 221320_at | 0,019246711 | 0,753543875 | 0,881541703 |
| **BCL2L11** | 225606_at | 0,000221674 | 0,998498713 | 0,999218832 | -0,019894506 | 0,909652227 | 0,942932187 | 222343_at | 0,100322368 | 0,170368769 | 0,517658952 |
| **BCL2L12** | 233110_s_at | 0,525174038 | 2,35431E-05 | 0,000168726 | 0,299044467 | 0,128702566 | 0,237819959 | | | | |
| **BCL2L13** | 217955_at | -0,802140896 | 7,52541E-12 | 1,61796E-10 | -0,693601629 | 5,56979E-06 | 7,94728E-05 | 217955_at | -0,121638158 | 0,024829978 | 0,130771217 |
| **BCL2L14** | 234191_at | 0,079435674 | 0,029168049 | 0,076013704 | 1,485196128 | 0,027470415 | 0,068676038 | 221241_s_at | 0,045904605 | 0,398088908 | 0,768184772 |
| **BCL2L2** | 209311_at | -0,012352437 | 0,886808726 | 0,95331938 | -0,101025874 | 0,384150815 | 0,562979642 | 209311_at | -0,076615132 | 0,11122014 | 0,382017004 |
| **BID** | 227143_s_at | 0,127456828 | 0,232044288 | 0,383765553 | 0,127636507 | 0,484786726 | 0,624346541 | 211725_s_at | 0,083764803 | 0,330444157 | 0,70554293 |
| **BIK** | 205780_at | 0,418701091 | 0,042348859 | 0,095842154 | -0,155669391 | 0,593477901 | 0,710501713 | 205780_at | 0,059680921 | 0,518129144 | 0,803854064 |
| **BIRC2** | 202076_at | -0,118517337 | 0,061113054 | 0,131393067 | 0,130485884 | 0,410890155 | 0,571533677 | 202076_at | -0,203258224 | 0,154073031 | 0,486870778 |
| **BIRC3** | 230499_at | 0,097690168 | 0,519492437 | 0,698067962 | -0,291564123 | 0,512621923 | 0,640777404 | 210538_s_at | 0,078011513 | 0,58636275 | 0,803854064 |
| **BIRC5** | 202095_s_at | -0,033889789 | 0,87065599 | 0,947802723 | -0,491424894 | 0,045786949 | 0,103270616 | 202095_s_at | -0,083759868 | 0,426872478 | 0,768184772 |
| **BMF** | 226530_at | -0,535088236 | 5,58182E-09 | 9,60074E-08 | -0,970864443 | 2,93757E-06 | 6,24234E-05 | | | | |
| **BNIP3** | 201849_at | -1,140219193 | 1,49595E-07 | 2,14419E-06 | -0,666187075 | 0,005656499 | 0,018492402 | 201849_at | -1,073878289 | 9,69135E-07 | 2,16184E-05 |
| **BNIP3L** | 221478_at | 0,410890486 | 0,000611366 | 0,00309279 | 0,42292523 | 0,05416939 | 0,115109953 | 221479_s_at | 0,411524671 | 0,003810322 | 0,025084619 |
| **BOK** | 223349_s_at | -0,320311433 | 0,074572337 | 0,152695737 | -0,875846261 | 0,000147414 | 0,001139109 | | | | |
| **CALR** | 212953_x_at | 0,203514368 | 0,036550432 | 0,088880035 | 0,379784659 | 0,006121438 | 0,019271194 | 212953_x_at | -0,006901316 | 0,959845793 | 0,984776852 |
| **CASP1** | 209970_x_at | 0,077639478 | 0,693474261 | 0,805929547 | 0,815552295 | 0,003594301 | 0,01454836 | 211366_x_at | 0,022534539 | 0,657254226 | 0,824175934 |
| **CASP10** | 210955_at | 0,057399741 | 0,245604641 | 0,391148131 | 0,146932264 | 0,574343903 | 0,707525098 | 211888_x_at | 0,037726974 | 0,466745178 | 0,768184772 |
| **CASP12** | 1564736_a_at | -0,055664869 | 0,06442175 | 0,135128548 | | | | | | | |
| **CASP14** | 231722_at | 0,0159423 | 0,727392612 | 0,834076862 | 0,216724479 | 0,416883388 | 0,571533677 | | | | |
| **CASP2** | 226032_at | 0,353495004 | 0,000226926 | 0,001301041 | 0,184686958 | 0,154712849 | 0,279799833 | 209811_at | 0,112379934 | 0,092068982 | 0,377527855 |
| **CASP3** | 202763_at | 0,318945366 | 0,002503351 | 0,009785827 | 0,605202906 | 0,00010845 | 0,000921825 | 202763_at | 0,078041118 | 0,435019544 | 0,768184772 |
| **CASP4** | 209310_s_at | 0,268913007 | 0,193275089 | 0,332433152 | 0,931618123 | 0,001624367 | 0,007266903 | 209310_s_at | -0,027955592 | 0,613445116 | 0,803854064 |
| **CASP5** | 207500_at | -3,63016E-05 | 0,999218832 | 0,999218832 | 2,342978126 | 4,5038E-05 | 0,000466074 | 207500_at | 0,040370066 | 0,603798985 | 0,803854064 |
| **CASP6** | 242323_at | -0,794385266 | 2,19757E-13 | 6,2997E-12 | 0,349752386 | 0,010678247 | 0,031298311 | 209790_s_at | 0,108868421 | 0,212025458 | 0,540322942 |
| **CASP7** | 207181_s_at | 0,169046825 | 0,271969957 | 0,425262114 | 0,188792727 | 0,240726027 | 0,393494468 | 207181_s_at | 0,026039474 | 0,620697442 | 0,803854064 |
| **CASP8** | 213373_s_at | 0,315449039 | 0,121605918 | 0,23240242 | 0,163727538 | 0,001453491 | 0,006863705 | 207686_s_at | 0,032162829 | 0,601978614 | 0,803854064 |
| **CASP9** | 203984_s_at | -0,397016241 | 0,002220652 | 0,009094097 | -0,649517433 | 0,004834404 | 0,017866275 | 210775_x_at | -0,186973684 | 0,00774441 | 0,007647606 |
| **CFLAR** | 210563_x_at | 0,224669048 | 0,278065851 | 0,4270297 | 0,464453438 | 0,081048576 | 0,153091755 | 210563_x_at | -0,407021382 | 1,48636E-06 | 2,34845E-05 |
| **CRADD** | 209833_at | -0,218166286 | 0,003104106 | 0,011123047 | -0,29055587 | 0,057550186 | 0,117625549 | 209833_at | -0,003175987 | 0,939637653 | 0,984776852 |
| **CSE1L** | 201112_s_at | 0,086887843 | 0,410915585 | 0,569979683 | -0,056046097 | 0,727906121 | 0,806562249 | 201112_s_at | -0,008342105 | 0,942844016 | 0,984776852 |
| **CYCS** | 208905_at | 0,235587128 | 0,019432636 | 0,052225209 | 0,059740574 | 0,740139476 | 0,806562249 | 208905_at | -0,014947368 | 0,921612391 | 0,984776852 |
| **CYLD** | 213295_at | -0,431975644 | 0,017204478 | 0,047728552 | -0,249669174 | 0,252414974 | 0,404816469 | 221905_at | 0,016947368 | 0,803566921 | 0,906882668 |
| **DIABLO** | 219350_s_at | 0,187546116 | 0,006096073 | 0,018985243 | -0,19834757 | 0,233077747 | 0,388462912 | 219350_s_at | 0,021064145 | 0,75879539 | 0,881541703 |
| **ENDOG** | 204824_at | -1,167643994 | 1,68537E-16 | 1,44942E-14 | -1,210524958 | 5,34947E-07 | 2,27353E-05 | 204824_at | -0,852266447 | 4,22038E-12 | 1,66705E-10 |
| **FADD** | 202535_at | 0,381041775 | 1,67088E-05 | 0,000130633 | 0,228165542 | 0,205375612 | 0,362798107 | 202535_at | 0,15590625 | 0,009958382 | 0,060516324 |
| **FAS** | 204780_s_at | 0,850944887 | 0,000165827 | 0,001018651 | 1,733523203 | 9,72059E-11 | 8,2625E-09 | 204781_s_at | 0,008253289 | 0,919594074 | 0,984776852 |
| **FASLG** | 210865_at | -0,067312885 | 0,148375914 | 0,271496353 | -0,236531505 | 0,410779801 | 0,571533677 | 210865_at | 0,179144737 | 0,128296105 | 0,422308013 |
| **GAPDH** | 217398_x_at | -0,004848182 | 0,943116021 | 0,976459263 | -0,194610682 | 0,271437498 | 0,427262728 | 212581_x_at | -0,188444079 | 0,095576672 | 0,377527855 |
| **GZMB** | 210164_at | 0,023087294 | 0,918391585 | 0,969585208 | 0,039023825 | 0,893735825 | 0,937870928 | 210164_at | 0,050861842 | 0,576261915 | 0,803854064 |
| **HRK** | 237187_at | 0,179454492 | 0,037205596 | 0,088880035 | 0,22410302 | 0,467174645 | 0,624346541 | 206864_s_at | 0,040539474 | 0,422235931 | 0,768184772 |
| **HTRA2** | 203089_s_at | 0,183396863 | 0,039205409 | 0,091126087 | 0,008679492 | 0,942994338 | 0,954220462 | 211152_s_at | 0,068276316 | 0,10580052 | 0,382017004 |
| **ICAM3** | 204949_at | 0,381612988 | 0,086416031 | 0,172832062 | 0,742059003 | 0,928296629 | 0,950665223 | 204949_at | 0,232190789 | 0,011055591 | 0,062385123 |
| **LRP1** | 200784_s_at | 0,440259235 | 0,000103657 | 0,000685729 | 1,261074554 | 5,60985E-06 | 7,94728E-05 | 200784_s_at | 0,056685855 | 0,326493364 | 0,70554293 |
| **MALT1** | 210018_x_at | 0,929538893 | 5,19515E-06 | 4,46783E-05 | 0,339975667 | 0,024280394 | 0,062540409 | 210018_x_at | 0,258243421 | 0,036545875 | 0,180445257 |
| **MAPK8** | 226048_at | -0,148775182 | 0,135828721 | 0,253940652 | 0,050110732 | 0,646803043 | 0,75312683 | 210477_x_at | 0,097169408 | 0,182417594 | 0,533740369 |
| **MCL1** | 200797_s_at | 0,00936331 | 0,924488222 | 0,969585208 | 0,581431743 | 0,003967412 | 0,015328636 | 200797_s_at | -0,084636513 | 0,573159569 | 0,803854064 |
| **MOAP1** | 212508_at | -0,528788286 | 2,75387E-06 | 2,96042E-05 | -0,551587048 | 0,000967245 | 0,005138486 | 212508_at | -0,639414474 | 1,81885E-05 | 0,000239482 |
| **NAIP** | 204860_s_at | -0,529953129 | 0,004229119 | 0,014548169 | 0,709220778 | 0,707316425 | 0,806562249 | 204861_s_at | -0,538794408 | 0,000362109 | 0,004086658 |
| **NLRP1** | 210113_s_at | -0,313767286 | 0,006181242 | 0,018985243 | -0,89421648 | 0,000594439 | 0,003886716 | 210113_s_at | 0,074861842 | 0,195557416 | 0,540322942 |
| **NLRP3** | 207075_at | -0,05598999 | 0,640009292 | 0,775222522 | 1,428877016 | 0,052028789 | 0,113396079 | 216016_at | 0,048230263 | 0,464703171 | 0,768184772 |
| **OPA1** | 212213_x_at | -0,15124291 | 0,060047116 | 0,131393067 | -0,289038193 | 0,019586084 | 0,053703779 | 212213_x_at | -0,051856908 | 0,223825802 | 0,552569949 |
| **PIDD1** | 219019_at | 0,12922441 | 0,109968916 | 0,214939246 | 0,260939766 | 0,217807128 | 0,370272118 | 219019_at | 0,210143092 | 0,000969969 | 0,008514171 |
| **PMAIP1** | 204285_s_at | 1,279502069 | 3,99679E-06 | 3,81915E-05 | 0,761317234 | 0,00538922 | 0,018323348 | 204285_s_at | 0,497213816 | 0,003250997 | 0,023348069 |
| **PYCARD** | 221666_s_at | 0,178078027 | 0,320620084 | 0,483742583 | 0,659493139 | 0,021500386 | 0,0571104 | 221666_s_at | 0,031855263 | 0,615236367 | 0,803854064 |
| **RIPK1** | 226551_at | 0,051317101 | 0,37395538 | 0,545087504 | -0,008187203 | 0,957827992 | 0,957827992 | 209941_at | -0,014182566 | 0,697294663 | 0,860723099 |
| **RIPK3** | 228139_at | 0,088335551 | 0,211327618 | 0,356356376 | 0,197590851 | 0,470687655 | 0,624346541 | | | | |
| **SH3GLB1** | 209090_s_at | -0,883493843 | 2,79819E-14 | 1,20322E-12 | -0,623139627 | 1,36059E-06 | 3,855E-05 | 209090_s_at | -0,871978618 | 1,18117E-13 | 9,33122E-12 |
| **TNF** | 1563357_at | -0,21065658 | 0,368162974 | 0,545087504 | 0,33052834 | 0,585573501 | 0,710501713 | 207113_s_at | 0,045916118 | 0,551660625 | 0,803854064 |
| **TNFRSF10A** | 231775_at | 0,05961151 | 0,615187035 | 0,755801215 | 0,307921076 | 0,299505491 | 0,462872122 | | | | |
| **TNFRSF10B** | 209295_at | 0,475853769 | 0,003080212 | 0,011123047 | 0,78287577 | 4,9349E-05 | 0,000466074 | 209295_at | 0,470712171 | 0,001533599 | 0,012115431 |
| **TNFRSF10C** | 210483_at | 0,118530016 | 0,031722055 | 0,080238139 | 0,264442206 | 0,346113364 | 0,516133963 | 206222_at | 0,028532895 | 0,64557678 | 0,822589769 |
| **TNFRSF10D** | 227345_at | 0,061254931 | 0,769286711 | 0,848187912 | 0,600106837 | 0,058120859 | 0,117625549 | 210654_at | 0,083440789 | 0,202389453 | 0,540322942 |
| **TNFRSF1A** | 207643_s_at | 0,819736779 | 1,51246E-06 | 1,85816E-05 | 1,042544123 | 1,83792E-05 | 0,000223176 | 207643_s_at | 0,438730263 | 1,0946E-06 | 2,16184E-05 |
| **TNFRSF1B** | 203508_at | 0,068769312 | 0,671562779 | 0,791156151 | 0,581959421 | 0,042354888 | 0,100004596 | 203508_at | 0,053345395 | 0,450184898 | 0,768184772 |
| **TNFSF10** | 202688_at | -0,19086215 | 0,56091882 | 0,719985351 | 0,102076765 | 0,716780952 | 0,806562249 | 202688_at | 0,139800987 | 0,38616479 | 0,768184772 |
| **TP53** | 201746_at | 0,489508727 | 0,002003913 | 0,008669539 | 0,400187397 | 0,000822379 | 0,004660146 | 201746_at | 0,185552632 | 0,266650646 | 0,619570619 |
| **TRADD** | 205641_s_at | -0,062418923 | 0,407251529 | 0,569979683 | -0,117700136 | 0,480314064 | 0,624346541 | 205641_s_at | 0,084902961 | 0,406371286 | 0,768184772 |
| **TRAF1** | 205599_at | 0,06684301 | 0,436149934 | 0,595379275 | 0,129438057 | 0,620882706 | 0,732986528 | 205599_at | 0,036384868 | 0,484601414 | 0,781296158 |
| **TRAF2** | 204413_at | 0,014563008 | 0,742026391 | 0,839661442 | -0,54873499 | 0,000661193 | 0,004014387 | 204413_at | -0,029050987 | 0,580605813 | 0,803854064 |
| **XAF1** | 228617_at | 0,073830039 | 0,763230223 | 0,848187912 | -0,091306709 | 0,731724778 | 0,806562249 | 206133_at | 0,154894737 | 0,050849629 | 0,236301217 |
| **XIAP** | 228363_at | -0,236210837 | 0,011043014 | 0,031656639 | 0,074200517 | 0,500182625 | 0,634560046 | 206537_at | 0,04521875 | 0,409767818 | 0,768184772 |

**Supplementary table 3. Evaluation of the optimal media for 3D RMS-O and 2D cultures of FNRMS and patients' clinical characteristics.**

| Sample name | Age (years) | Sex | FN-RMS Histological | Status | Sampling location | Location of primary tumor (in case of | Sampling type | Tumor treatment | Interval to last treatment | Culture in 3D M3 | Culture in 2D DMEM | Culture in 2D in complex RMS-O |
|---|---|---|---|---|---|---|---|---|---|---|---|---|
| RMS1 | 8 | M | Embryonal | **Relapse** | Parameningeal | NA | Biopsy | Yes | 3 weeks | OPTIMAL | NA | NA |
| RMS2 | 7 | M | Embryonal | **Relapse/metastatic** | Lung | Ischioanal | Resection | Yes | On-going | OPTIMAL | NA | NA |
| RMS10 | 43 | M | Pleomorphic | **Relapse/metastatic** | Gluteal | ND | Resection | Yes | On-going | OPTIMAL | NA | NA |
| RMS11 | 31 | F | Embryonal | **Relapse/metastatic** | Thoracic | Cervical (thyroid) | Resection | Yes | On-going | OPTIMAL | NA | NA |
| PDXA | 5 | M | Embryonal | **Relapse** | Abdomen | NA | ND | Yes | ND | OPTIMAL | NA | NA |
| PDXC | 6 | M | Embryonal | **Relapse** | Gluteal | Gluteal | Biopsy | No | NA | OPTIMAL | NA | NA |
| RMS3 | 1 | M | Embryonal | **Primary** | Gluteal | NA | Biopsy | No | NA | NON-OPTIMAL | NA | NA |
| RMS6 | 4 | F | Embryonal | **Primary** | Pelvis | NA | Resection | Yes | 1 month | NON-OPTIMAL | NA | NA |
| PDXB | 5 | F | Embryonal | **Primary** | Parapharyngeal | NA | ND | No | NA | NON-OPTIMAL | NA | NA |
| RMS5 | 1 | M | Embryonal | **Primary** | Gluteal | NA | Resection | Yes | Few days | NON-OPTIMAL | OPTIMAL | OPTIMAL |
| RMS4 | 7 | M | Embryonal | **Primary** | Paratesticular | NA | Biopsy | No | NA | NON-OPTIMAL | NON-OPTIMAL | OPTIMAL |
| RMS12 | < 1 | M | Embryonal | **Primary** | Foot | NA | Resection | Yes | ND | NON-OPTIMAL | NON-OPTIMAL | OPTIMAL |
| RMS7 | 1 | F | Embryonal | **Primary** | Thoracic | NA | Biopsy | No | NA | NA | NON-OPTIMAL | OPTIMAL |
| RMS8 | 21 | F | Spindle cell | **Primary** | Infratemporal | NA | Resection | No | NA | NA | NON-OPTIMAL | OPTIMAL |
| RMS9 | 23 | M | Spindle cell | **Primary** | Intermaxillary | NA | Resection | No | NA | NA | NON-OPTIMAL | OPTIMAL |

**M3 culture condition was defined as optimal based on its suitability to sustain continuous organoid growth. For 2D cultures, RMS-O like and DMEM culture conditions were defined as optimal or non-optimal based on their suitability to sustain continuous proliferation of cells and to support cell growth after a freezing/thawing cycle. NA = Not Applicable. ND = Not Documented.**
